# Multi-omic Characterization of HIV Effects at Single Cell Level across Human Brain Regions

**DOI:** 10.1101/2025.02.05.636707

**Authors:** Junchen Yang, Kriti Agrawal, Jay Stanley, Ruiqi Li, Nicholas Jacobs, Haowei Wang, Chang Lu, Rihao Qu, Declan Clarke, Yuhang Chen, Yunzhe Jiang, Donglu Bai, Suchen Zheng, Howard Fox, Ya-chi Ho, Anita Huttner, Mark Gerstein, Yuval Kluger, Le Zhang, Serena Spudich

## Abstract

HIV infection exerts profound and long-lasting neurodegenerative effects on the central nervous system (CNS) that can persist despite antiretroviral therapy (ART). Here, we used single-nucleus multiome sequencing to map the transcriptomic and epigenetic landscapes of postmortem human brains from 13 healthy individuals and 20 individuals with HIV who have a history of treatment with ART. Our study spanned three distinct regions—the prefrontal cortex, insular cortex, and ventral striatum—enabling a comprehensive exploration of region-specific and cross-regional perturbations. We found widespread and persistent HIV-associated transcriptional and epigenetic alterations across multiple cell types. Detailed analyses of microglia revealed state changes marked by immune activation and metabolic dysregulation, while integrative multiomic profiling of astrocytes identified multiple subpopulations, including a reactive subpopulation unique to HIV-infected brains. These findings suggest that cells from people with HIV exhibit molecular shifts that may underlie ongoing neuroinflammation and CNS dysfunction. Furthermore, cell–cell communication analyses uncovered dysregulated and pro-inflammatory interactions among glial populations, underscoring the multifaceted and enduring impact of HIV on the brain milieu. Collectively, our comprehensive atlas of HIV-associated brain changes reveals distinct glial cell states with signatures of pro-inflammatory signaling and metabolic dysregulation, providing a framework for developing targeted therapies for HIV-associated neurological dysfunction.

## 1 Introduction

HIV infection is known to have pervasive effects on the central nervous system (CNS) [1]. However, little is known about the localized effects of HIV on different brain regions or on distinct cell types within the brain. Despite anti-retroviral therapy (ART), some people with HIV still experience chronic and sometimes progressive neurologic dysfunction, and it is unclear whether the underlying causes of these changes can be directly attributed to viral persistence – whether viral particles, nucleic acid, or proteins - or the inability of the brain to resolve HIV-mediated neuroinflammation and activation. Animal models have been helpful at providing insights into cellular changes that occur during HIV infection, but as HIV is specific to humans, animal studies either employ humanized models, engineered viruses, or naturally occurring species specific viruses such as simian immunodeficiency viruses that differ in natural history from HIV in the human host [2, 3]. There remains a lack of comprehensive human studies at the tissue level to confirm that phenomena observed in these conditions occur in humans in the setting of both a distinct virus and distinct immune and CNS cellular profiles. Recent in-depth studies that have been carried out using human-derived tissue focus on characterizing changes in specific cell types (such as microglia and astrocytes) and have helped uncover key mechanisms that may contribute to the detrimental effects of HIV in the brain [4, 5]. However, these studies lack a comprehensive view of cellular interactions in the brain.

As a member of the Single Cell Opioid Responses in the Context of HIV (SCORCH) consortium [6], we have been working to decipher the transcriptomic and epigenetic changes that occur in widespread regions and cell types in the brain as a result of HIV infection. In this study, we present a comprehensive overview of the lasting effects of HIV on the human brain using single nucleus multiomic sequencing (snRNA+snATAC) of postmortem brains from 13 people without HIV (CTR) and 20 people with HIV (PWH) who have a history of treatment with ART. Our study encompasses multiple brain regions – specifically, the prefrontal cortex (PFC), the insular cortex (INS), and the ventral striatum (VST). This study is among the first of its kind to sequence a large cohort of individuals, revealing widespread and persistent transcriptional and epigenetic changes in numerous cell types across brain regions. We generated a comprehensive cell atlas and identified consistent cross-regional effects of HIV infection across multiple cell types, along with distinct region-specific signals. We then thoroughly characterized the molecular signatures of microglia and astrocytes, unveiling their complex states marked by immune activation and metabolic dysfunction in the context of HIV infection and the subsequent host response. Besides pro-inflammatory signaling pathways, HIV microglia also exhibit unique signatures of hypoxia and lipid accumulation. By combining transcriptomic and epigenetic information, we identified eight subpopulations of astrocytes, with one population of reactive astrocytes that is unique to HIV. Cell-cell communication within glial populations revealed several pro-inflammatory and dys-regulated interactions, suggesting persistent changes to the brain. Together, these findings illustrate the extensive and enduring influence of HIV across brain regions, underscore the dynamic involvement of glial cells in chronic HIV infection, and highlight the necessity of continued investigation into these molecular changes.

## 2 Results

To characterize the effects of HIV at both a transcriptomic and a epigenetic level in the human brain, we profiled autopsy brain tissue donated by 20 PWH and 13 people without HIV (CTR) across PFC, INS, and VST using 10x Multiome (Methods **Section** 6.1). All of the PWH had a history of treatment with ART (approximately one third on recently suppressive ART) and while some of the individuals had records indicating cognitive or neurologic impairments in life, only one donor was determined on pathologic examination to have HIV encephalitis (HIVE). The groups did not differ by sex: the HIV group included 4 women and 16 men, while the CTR group comprised 3 women and 10 men. The HIV and CTR groups had a median age of 47.6 and 43 years old, respectively. Other detailed metadata, including details of HIV parameters in the PWH, are provided in supplementary table 1.

For the RNA modality, we processed each sample using Nextflow [7] with Seurat [8] as the core processing package (see Methods **Section** 6.4 for details). In total, 88 samples with 654, 918 nuclei from the RNA modality passed quality control and processing steps. Fig. 1b shows the t-SNE visualization of the filtered nuclei from the RNA modality colored by regions and conditions, separately, with the number of nuclei per condition in each brain region shown in Fig. 1c. For the ATAC modality, we performed peak calling for cell types defined using the RNA modality, and we processed each sample using signac [9]. In total, 86 samples with 458, 375 nuclei from the ATAC modality passed quality control and processing steps.

**Figure 1:**
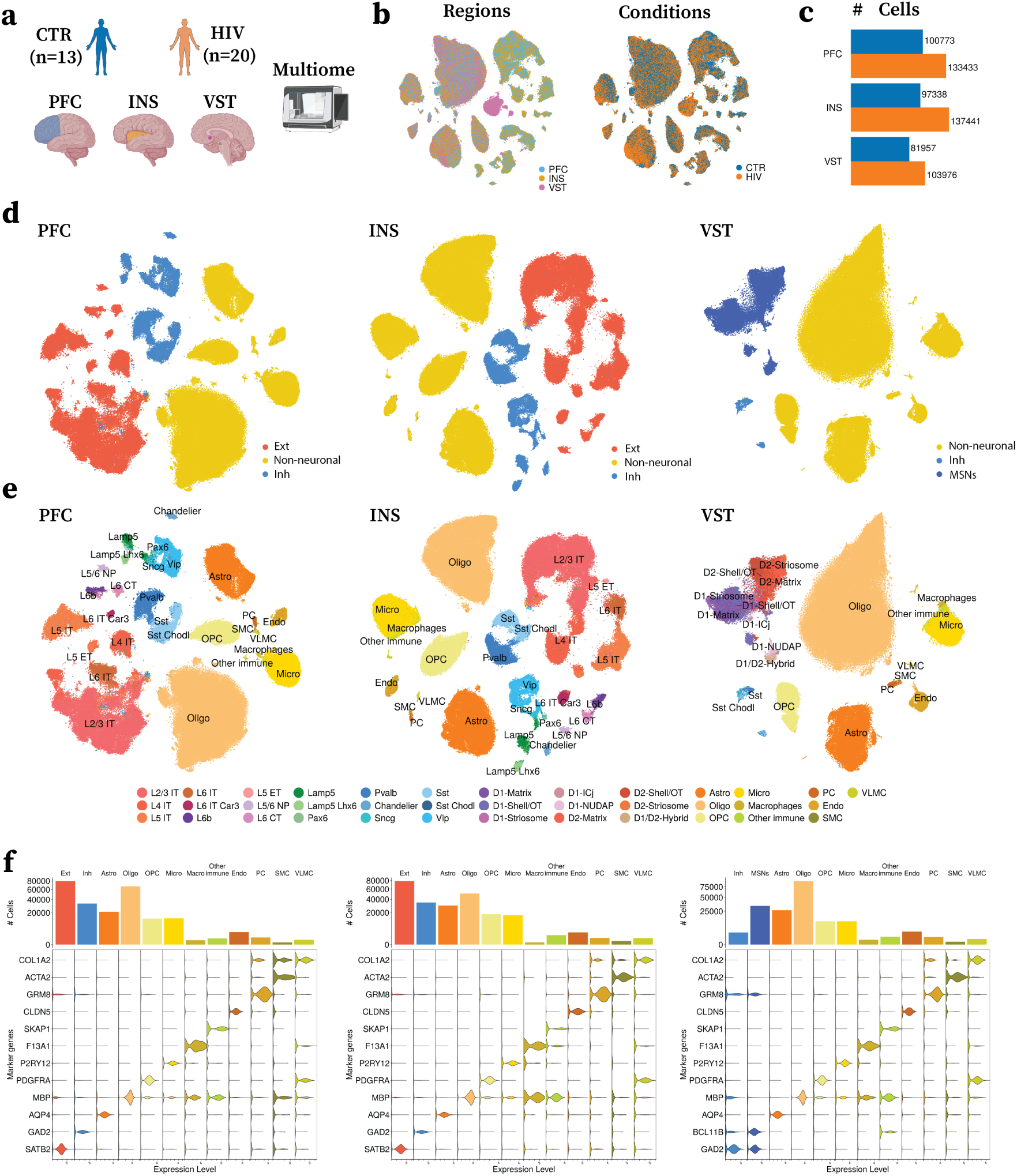
Single nuclei profiling of HIV effects across different brain regions. **a**: Overview of the study design. Three brain regions, including prefrontal cortex (PFC), insular cortex (INS), and ventral striatum (VST), are dissected and profiled using 10x Multiome of post-mortem tissue from 20 PWH and 13 CTR. **b**: t-SNE visualization of the cells in the RNA data after processing, colored by brain regions (left) and conditions (right), respectively. **c**: Total number of cells grouped by regions and conditions after processing. **d-e**: t-SNE visualization of the cells after cell annotations. Cells are labeled and colored by the annotated coarse-grained cell types (**d**) and fine-grained cell types (**e**) across regions. **f** : Cell numbers per cell type and marker gene validation of the annotations. Bar plot visualizations (top) show the number of cells per cell type in each region. The stacked violin plots (bottom) show the expression levels of the marker genes (rows) across the cell types (columns). Rows and columns are sorted such that the markers and the cell types are in the same order. Left-to-right, panels correspond to each brain region(PFC, INS, and VST, respectively).

### 2.1 Comprehensive annotation of cell types across regions

To efficiently and accurately annotate our data, we integrated external single-nucleus RNA-seq brain datasets from various regions to create a reference atlas to label the fine-grained cell subtypes in the RNA modality using multiple references from [10, 11, 12, 13] (Methods **Section** 6.4). As illustrated by the t-SNE visualizations in Fig. 1d and e, we identified nine glutamatergic excitatory neuron (Ext) types and ten GABAergic inhibitory neuron (Inh) types in both the PFC and INS. Specifically, the Ext subtypes include L2/3 IT, L4 IT, L5 IT, L6 IT, L6 IT Car3, L6b, L5 ET, L5/6 NP, and L6 CT, while the Inh subtypes comprise Lamp5+, Lamp5+Lhx6+, Pax6+, Pvalb+, Chandelier+, Sncg+, Sst+, Sst+Chodl+, and Vip+ Inh. For VST, we identified nine different subgroups of medium spiny neurons (MSNs) with specific expression of dopamine receptor 1 (D1) and dopamine receptor 2 (D2), namely, D1-Matrix, D1-Shell/OT, D1-Striosome, D1-ICj, D1-NUDAP, D2-Matrix, D2-Shell/OT, D2-Striosome, and D1/D2-Hybrid MSNs. In addition, we identified four glial subtypes in all regions, including Astrocytes (Astro), Oligodendrocytes (Oligo), Oligodendrocyte Precursor Cells (OPC), and Microglia (Micro). We also found populations of neurovascular cells, such as Pericytes (PC), Endothelial cells (Endo), Smooth muscle cells (SMC), Vascular leptomeningeal cells (VLMC), Macrophages, and other peripheral immune cells, including T cells and myeloid lineage cells (Fig. S1). We further validated the quality of the annotations using canonical marker genes for each cell type using external datasets (Fig. 1e, Fig. S2-Fig. S4).

Next, we examined the quality of our annotations on the ATAC modality, as shown in the ATAC t-SNE colored by the RNA modality based cell annotations (Fig. 2a). For each nucleus, we quantified its neighborhood purity by calculating the percentage of its 100 nearest neighbors that share the same cell type label. On average, the neighborhood purity is 90% for PFC, 87% for INS, and 83% for VST, suggesting that cells of the same type tend to cluster together in the ATAC space and that the RNA and ATAC annotations are generally consistent. To quantify the open chromatin profiles at the gene level, we also aggregated the fragments within the gene body and its promoter region, and we computed the gene activity score for each gene. We examined the specificity of some top marker genes, namely, *SATB2* for Ext, *GAD2* for Inh, *BCL11B* for MSNs, *PDGFRA* for OPC, and *P2RY12* for microglia (Fig. 2b). In addition, we identified the top 50 most active genes for each cell type in each region in the ATAC modality. Most genes show strong specificity for the corresponding cell type (Fig. 2d) – e.g., different subtypes of excitatory and inhibitory neurons. In contrast, some cell types, such as different subtypes of medium spiny neurons, exhibit less distinct open chromatin signatures, suggesting that their epigenetic profiles are more uniform when compared to their transcriptomic characteristics.

**Figure 2:**
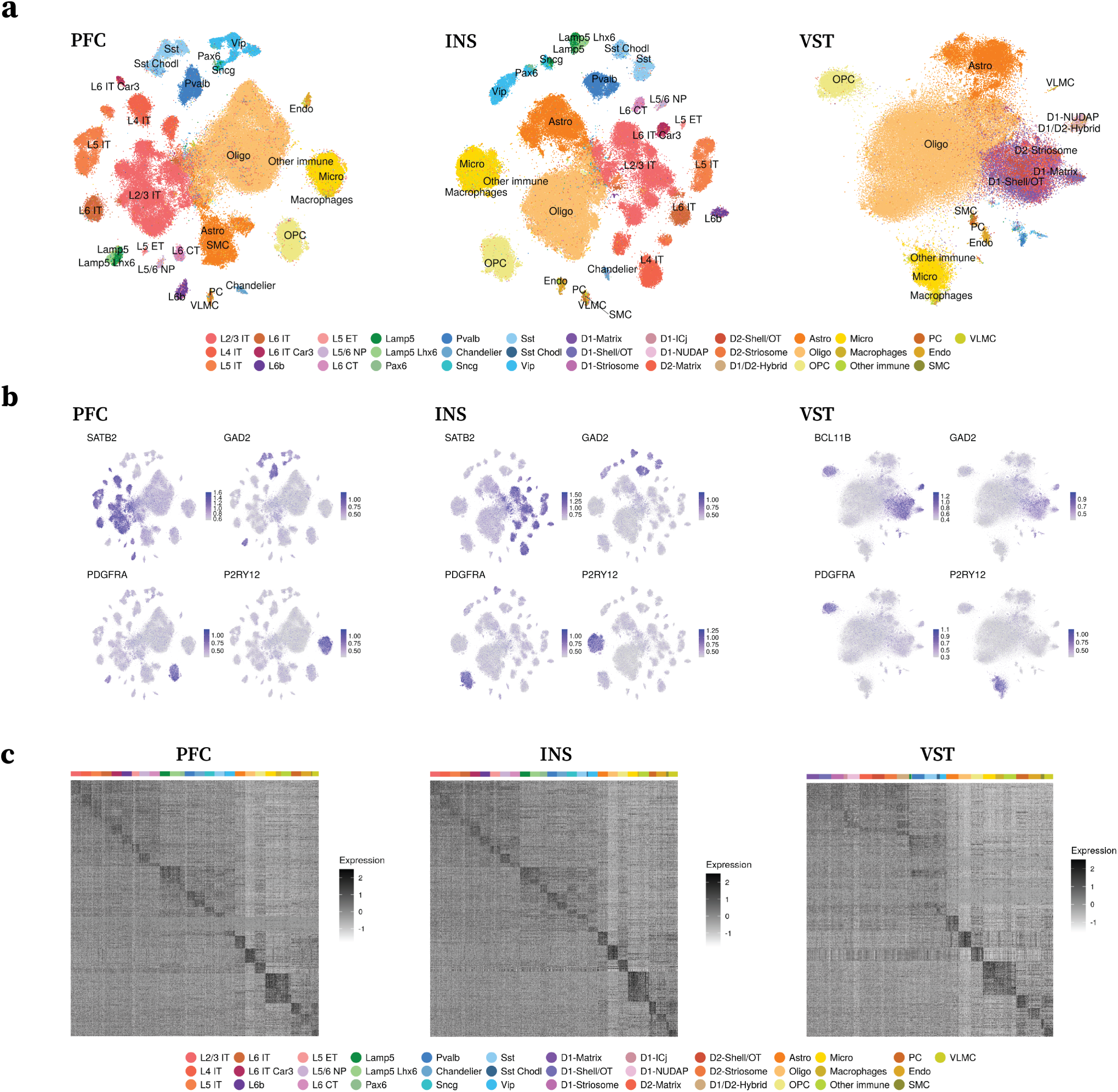
Epigenetic characterization of the brain regions. **a**: t-SNE visualization of the ATAC data for each brain region (PFC, INS, and VST), colored by the cell annotations. **b**: t-SNE visualization of the gene activities of marker genes in each region. SATB2 - Excitatory neurons, GAD2 - Inhibitory neurons, BCL11B - Medium spiny neurons, PDGFRA - OPC, P2RY12 - Microglia. **c**: Heatmap visualization of the top active genes (y-axis) across different cell types (axis) for each region. The values are log-normalized gene activity scores.

We also performed viral mapping to detect HIV DNA and RNA reads across samples in the ATAC and RNA (see Method **Section** 6.12 for details). After examining the mapped barcodes passed QC, we found that the detected DNA and RNA reads are enriched in the PFC region of the HIVE individual. Specifically, two microglia and two myeloid cells are identified with HIV RNA, whereas two microglia, one myeloid cell, and one astrocyte are identified with HIV DNA. In addition, one microglia and one myeloid cell are identified with both HIV DNA and RNA.

### 2.2 Cross-region differential analysis reveals widespread HIV effect

Following the cell annotations, we first examined the transcriptomic differences between PWH and CTR in each cell type for each region. Differential gene expression analysis using EdgeR [14, 15] revealed a large number of differentially expressed genes (DEGs) in multiple neuronal and glial cell types across regions (Fig. 3a). Fewer DEGs were observed in VST than in PFC and INS, likely due to the fact that fewer VST samples limit the overall statistical power. On the other hand, both PFC and INS exhibit strong DE signals in cell types such as L2/3 IT excitatory neurons, Vip+ inhibitory neurons, astrocytes, microglia, and endothelial cells.

**Figure 3:**
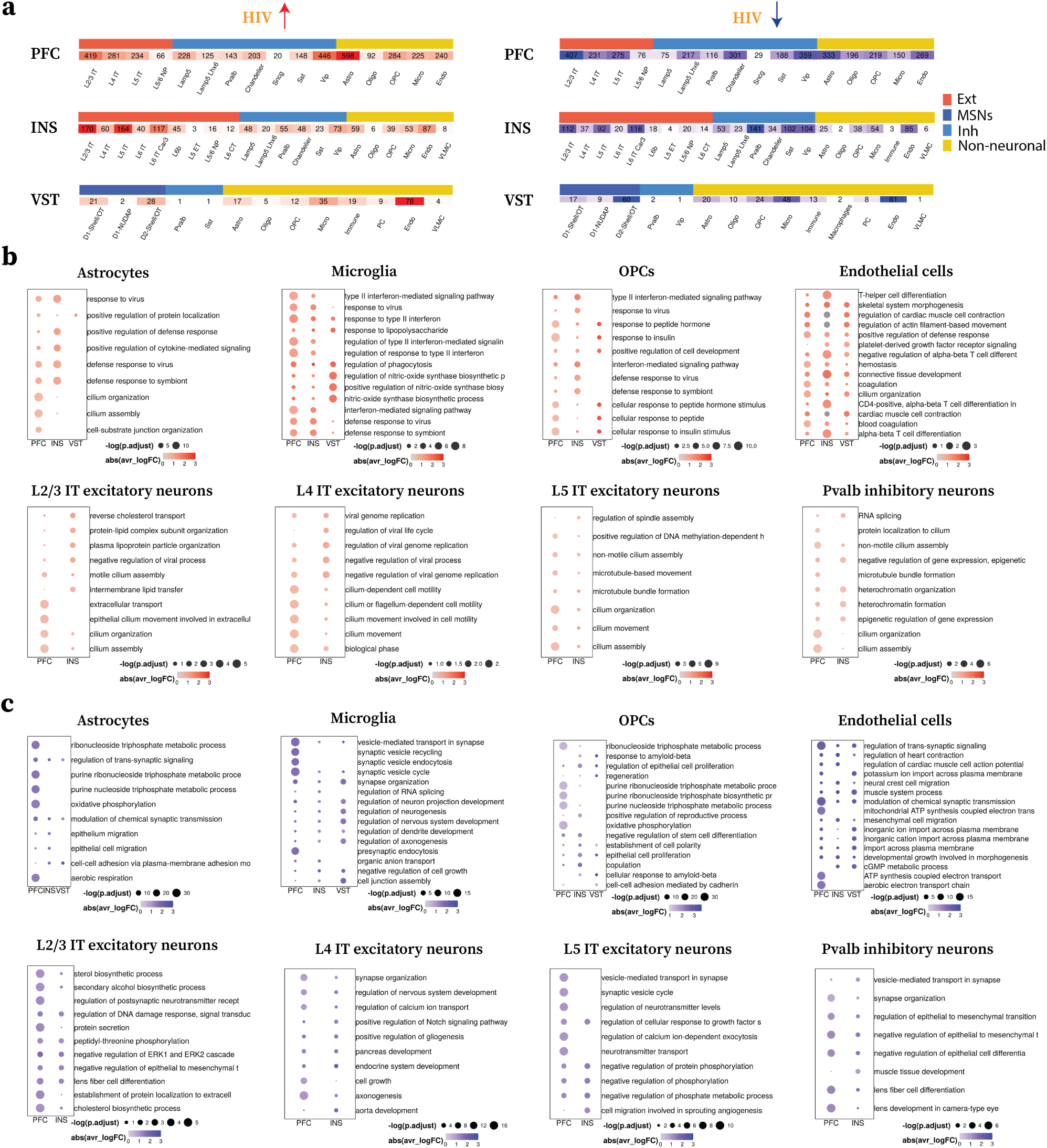
Differential gene expression analysis reveals strong signals associated with HIV. **a**: Total number of differentially expressed genes (DEGs) in each cell type that are upregulated in HIV (left) and downregulated in HIV (right). The heatmap is splited by regions and colored by the coarse-grained cell types (Ext, MSNs, Inh, Glial). **b**: Dot plot visualizations of the top upregulated pathways in HIV from each cell type across regions. **c**: Dot plot visualizations of the top downregulated pathways in HIV from each cell type across regions.

The gene ontology (GO) analysis of these region-specific DEGs reveals many top pathways enriched in genes upregulated in PWH (Fig. 3b) and genes downregulated in PWH (Fig. 3c) for different cell types across regions. Among the pathways that are upregulated in PWH, several glial cell types exhibit multiple immune response pathways, such as cytokine and interferon signaling, suggesting widespread chronic inflammatory signals among these cells in PWH. On the other hand, we observed cell-type specific signatures as well. For example, the biosynthetic process of nitric oxide (NO) synthase and its regulation are among the top pathways in microglia. The inducible form of nitric-oxide synthase (iNOS), a hallmark of pro-inflammatory microglia, produces high levels of NO in response to pro-inflammatory and degenerative stimuli, and NO is one of the reactive oxygen species (ROS) associated with neuroinflammation and neurodegeneration [16, 17, 18, 19, 20]. Insulin response pathways are enriched in OPC across regions, suggesting possible compensatory mechanisms to counteract neuroinflammation-driven metabolic stress and impaired myelination [21, 22]. Interestingly, our analyses revealed that endothelial cells were enriched for pathways typically associated with T cell differentiation. This observation was primarily driven by elevated expression of several cytokine receptors—namely *IL6R*, *IL18R1*, *BCL6*, and *IL4R* (Fig. S5)—indicating a possible inflammatory response to the corresponding cytokines. We also observed significant enrichment of cilium organization and assembly pathways in multiple glial and neuronal cell types mostly within PFC. Given primary cilia have been found to play important roles in different neurological diseases [23, 24], these findings raise the possibility that chronic HIV-associated inflammation could disrupt ciliary structure and function and induce neuro-degeneration. As for the DEGs downregulated in people with HIV, multiple synaptic transmission and signaling-related pathways are enriched across glial and neuronal cell types in different regions, indicating their active participation in maintaining healthy synaptic function.

Next, we focus on cross-region comparison of the DEGs. We compared the effect size of the DEGs between each pair of regions (Fig. 4a-c, INS-PFC, PFC-VST, INS-VST). Many of the DEGs show co-upregulation or co-downregulation patterns across regions. Multiple genes related to immune response have strong signals across regions. For example, *CD163*, an anti-inflammatory marker, is upregulated across all regions in microglia; *STC1*, a gene that encodes protein stanniocalcin-1, has been identified to play a critical role in endothelial cells in regulating the migration of inflammatory cells and promoting angiogenesis [25, 26]. Besides the consistent cross-region DE signals, we also observed regional differences for certain DEGs. For instance, L4 IT excitatory neurons from INS show stronger expression of *B2M* compared to PFC. *B2M* is a component of the Major Histocompatibility Complex (MHC) class I molecule whose expression level is elevated in neurons following stress and other neurodegenerative conditions [27] (Fig. 4a). Compared to PFC, INS astrocytes show more elevated expression of *PDGFRB*, which encodes a cell surface receptor that binds to platelet-derived growth factors (PDGFs). One of the growth factors, PDGF-B, can be induced by HIV-1 and has been linked to the increased proliferation of astrocytes and release of pro-inflammatory cytokines [28].

**Figure 4:**
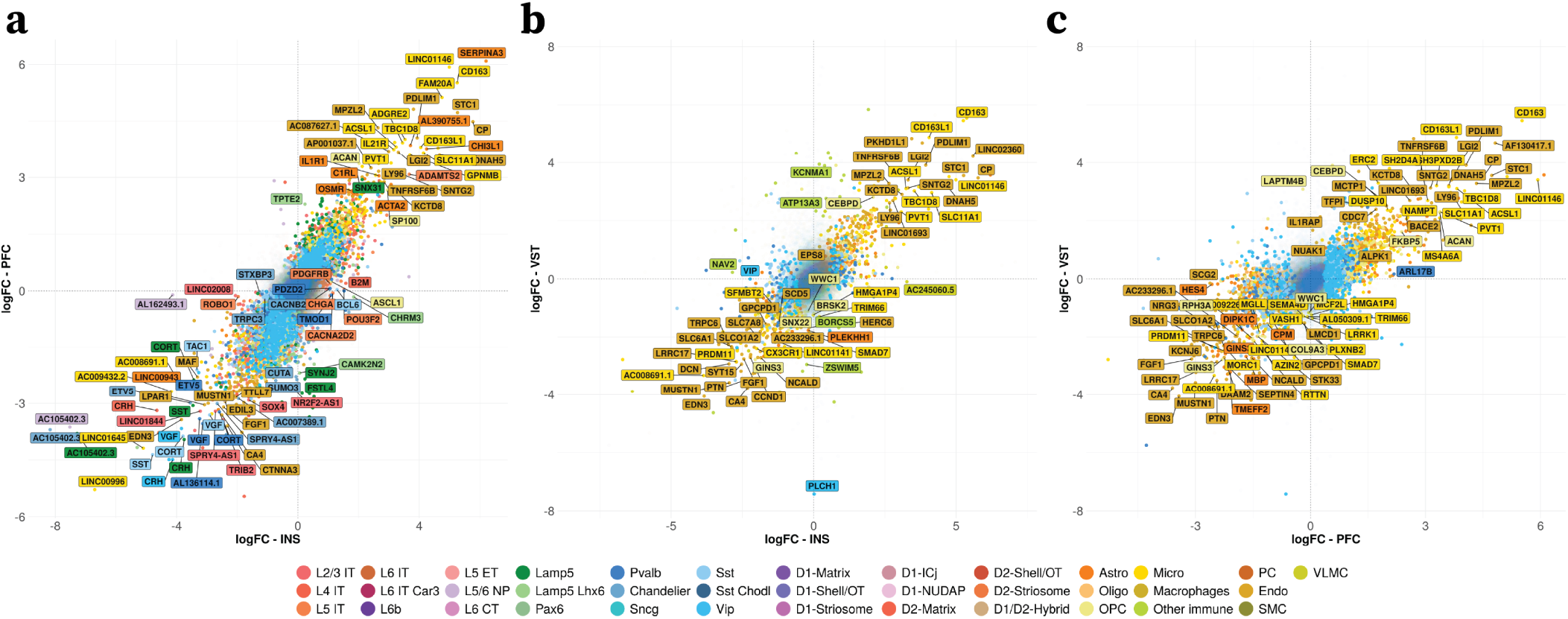
Cross-region comparison of differential signals. **a-c**: Comparison of DEGs and their effect size (logFC) between HIV and CTR for each pair of regions. Genes are colored by the corresponding cell types and the top DEGs are labeled with gene names. **a**: logFC of the DEGs from INS (x-axis) vs logFC of the DEGs from PFC (y-axis). **b**: logFC of the DEGs from INS (x-axis) vs logFC of the DEGs from VST (y-axis). **c**: logFC of the DEGs from PFC (x-axis) vs logFC of the DEGs from VST (y-axis).

To measure the concordance of DEGs between regions, we compute the Spearman correlation of log fold-change for each gene between regions for each cell type (Fig. S6a). Multiple glial and vascular cell types (e.g., endothelial cells, microglia, and astrocytes) exhibit strong concordance across all three regions, suggesting a consistently persistent HIV effect on these groups. On the other hand, INS and PFC show consistently higher concordance DEGs across cell types compared to VST, suggesting that the effect of HIV might be more similar within these two cortical regions.

Besides analyzing the differential signals for each region separately, we also performed the differential analysis jointly across regions. Specifically, we used the contrast model from EdgeR to identify the average effect of HIV across all three regions (see Methods **Section** 6.5 for details). GO analysis further verifies the pathways previously identified for each region, e.g., several immune response-related pathways for the DEGs that are upregulated in HIV across multiple glial cell types (Fig. S6b), and the pathways related to synaptic signaling and organization are enriched for these glial cell types in the downregulated DEGs (Fig. S6c).

In addition to our DEG analysis, we also performed differential analysis on the paired ATAC data. We first performed differential accessible peak analysis using EdgeR to identify peak regions that are differential between PWH and CTR in each cell type for each region. Several neuronal and glial cell types such as L4 IT excitatory neurons and astrocytes exhibit strong signals of differential accessible peaks (Fig. S8a). We also ran differential analysis at the gene level on the gene activity scores to identify differentially active genes. Astrocytes exhibit strong signals as well as consistent active genes between regions (Fig. S8b and c), suggesting widespread transcriptional regulation. Lastly, we combined results of DEG analysis with the results from the differential gene activity analysis to compare the differential signals between gene expression and gene activity for the top DEGs (Fig. S8d). Many genes showed consistent directions in terms of gene expressions and gene activities, such as *CD163* from PFC microglia and *SERPINA3* from PFC and INS, confirming the consistency between chromatin accessibility and gene expression. Together, these results of differential analysis provide a comprehensive overview of the widespread HIV effect across cell types and brain regions at both transcriptomic and epigenetic level.

### 2.3 Characterization of microglia states and corresponding molecular signatures

Microglia are thought to be reservoirs of HIV in the brain because they are actively infected by HIV and are a population of long-lived cells. [29]. Several studies, have suggested that microglia may be integrating HIV into their genome, allowing for possibility of latent HIV infection and causing a possible fundamental change in microglia function with HIV infection. Studies exploring the continued presence of HIV associated neurological diseases (HAND) despite ART therapy have found evidence that microglia may be playing a major role in the continued persistence of neurological damage after treatment for HIV [30]. Principal Component Analysis (PCA) reveals strong separations of microglia between PWH and CTR across all three regions (Fig. 5a). In line with the PCA results, many key DEGs exhibit consistent upregulation/downregulation patterns across regions (Fig. 5b). For example, a previous study has identified the upregulation of *IGF2R* in activated microglia may contribute to HIV persistence and expansion in the CNS, and may also control the induction of microglial inflammatory genes [31]. The solute carrier family 11 member 1 (*SLC11A1*) gene is associated with disrupted synaptic function and apoptosis, serving as a common marker of HIV-associated neurocognitive disorders (HAND) [32, 33]. *TLR2* has been identified to be upregulated by HIV gp-120 in microglia and leads to the production of inflammatory cytokines [34]. Moreover, many of the upregulated DEGs are enriched in pathways of defense response to virus, regulation of innate immune response, and interferon-mediated signaling pathways (Fig. 5c). Among them, guanylate binding protein 2, *GBP2*, participates in the regulation of cytokine production and pyroptosis and has been previously reported to be upregulated in the microglia of HIVE patients [5]. We also identified several signal transducer and activator of transcription (STAT) genes in these pathways, which are key transcription factors for the activation and inflammatory responses of microglia.

**Figure 5:**
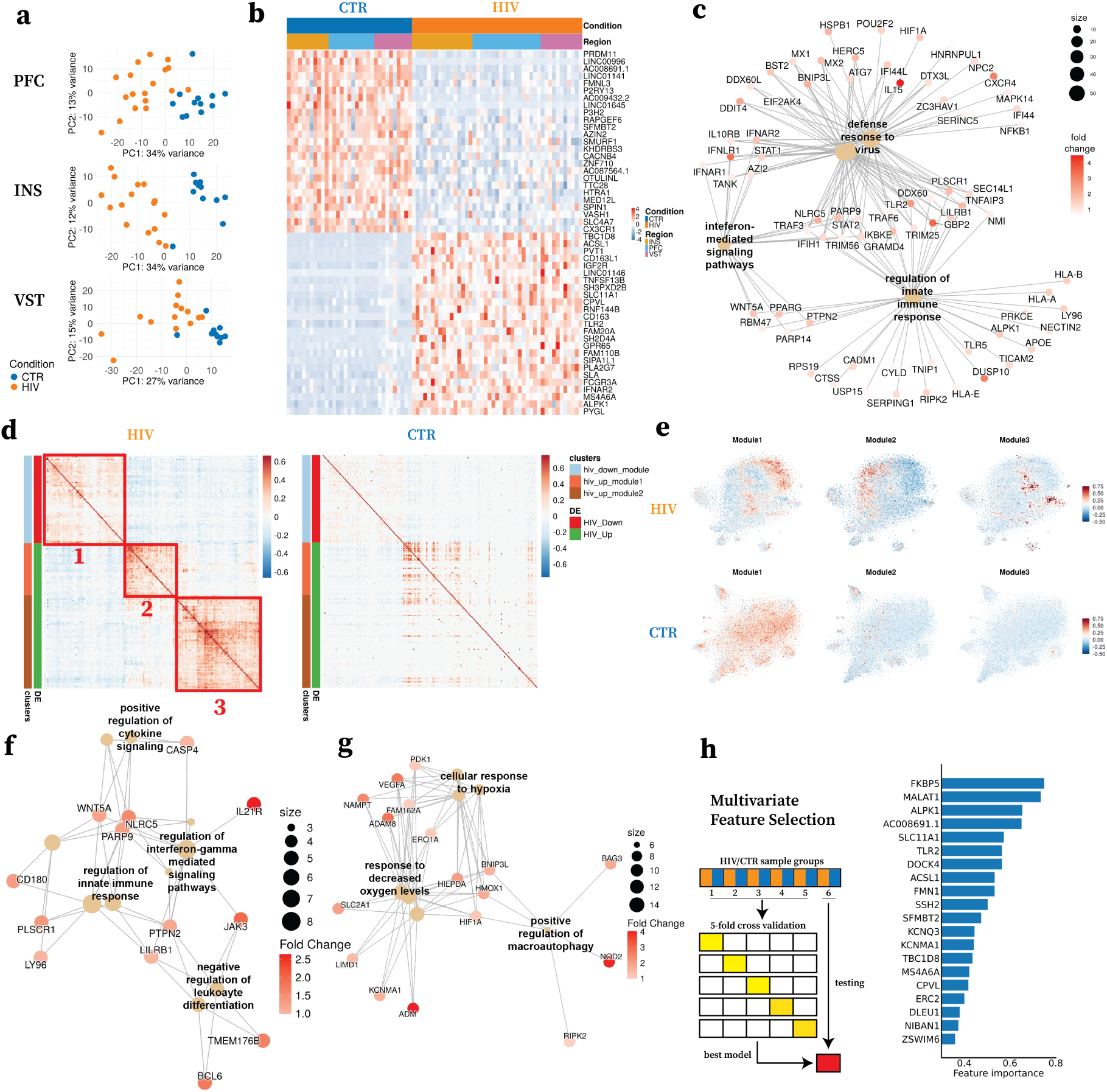
The effects of HIV on microglia of the INS, PFC, and VST. **a**: Principal Component Analysis (PCA) visualization of the microglia samples for each region. Each sample is colored by the corresponding condition label. **b**: Heatmap visualization of the top DEGs of the HIV main effect. Columns are different samples grouped by conditions and regions, and rows are the top DEGs with 25 each in the HIV-upregulated group and HIV-downregulated group. The expression levels inside the heatmap are z-scored for each gene. **c**: Visualization of top enriched GO pathways for the upregulated DEGs across regions. **d**: Heatmap visualization of the identified gene modules in HIV samples (Left) compared to the control samples (Right). Rows and columns are DEGs grouped by direction of the differential expression, and values are pair-wise correlation coefficients. Three gene modules are identified via hierarchical clustering (labeled as 1,2, 3). **e**: t-SNE visualization of HIV microglia (top row) and CTR microglia (bottom row) colored by the gene module scores from each of three identified gene modules (columns). **f** : Visualization of the top GO pathways associated with gene module 2. **g**: Visualization of the top GO pathways associated with gene module 3. **h**: Multivariate feature selection to select most informative genes predicting the conditions. Left: model training schemes. Right: Top selected genes (y-axis) and their feature importance (x-axis) for the best model.

We identified multiple markers that are reflective of different states of microglia, e.g., interferon response and stimulation markers such as *IL15*, *IFNAR1*, *IFNLR1*, *IFI44*, *IFI44L*, and genes associated with neuro-degenerative disorders in microglia such as *APOE* and *LPL* (Fig. S7). We also examined other marker genes of different microglia states reported from different literature, including homeostatic, anti-inflammatory, antigen presentation, and proliferative markers (Fig. S7) [35, 36, 37, 38, 39, 40]. Homeostatic markers *CX3CR1*, *P2RY12*, *SALL1* are significantly downregulated across regions (adjusted p-values = 5.28 *×* 10*^−^*^7^, 2.41 *×* 10*^−^*^5^, 5.6 *×* 10*^−^*^4^, respectively). On the other hand, we did not observe a significant upregulation of marker genes associated with antigen presentation and pro-liferative states at neither the transcriptomic nor epigenetic level. Although canonical anti-inflammatory cytokine marker genes such as *IL4*, *IL10*, and *IL13* do not exhibit significant differential expressions, we observe genes like *CD163* with significant upregulation at transcriptomic and epigenetic levels, indicating possible HIV context-specific anti-inflammatory signatures for microglia.

To further characterize the complex dynamic states of microglia, we examined the pairwise correlation patterns of differentially expressed genes using CS-CORE [41], a statistical approach for estimating cell-type-specific co-expressions that corrects for sequencing depth variations and measurement errors in scRNA-seq data. We identify three unique gene modules that correspond to three groups of microglia, with gene module 3 only appearing in PWH (Fig. 5d and e). Gene module 1 corresponds to a group of genes that are downregulated in HIV, including many marker genes that correspond to microglia homeostasis (supplementary table 2). On the contrary, gene modules 2 and 3 represent genes that are upregulated in PWH. Specifically, genes from module 2 are enriched in many pro-inflammatory pathways such as regulations of cytokines and interferon-gamma mediated signaling (Fig. 5f), whereas genes from module 3 are associated with many hypoxia related pathways (Fig. 5g). Interestingly, several genes linked to lipid accumulation—such as *ACSL1*, *NAMPT*, and *PLIN2* —are also found to be enriched in module 3, suggesting the concerted interplay between hypoxia and lipid metabolism. Abnormal accumulation of lipids and lipid droplets (LDs) – the lipid-storing organelles, can be triggered due to exposure to hypoxic conditions, and has been recognized as a hallmark for inflammation of myeloid cells including microglia [42, 43, 44, 45]. The corresponding dysfunctional state of microglia – LD-accumulating microglia (LDAM), has been characterized in the neuro-HIV context, where HIV-Transgenic rats show higher levels of LDs formation in hippocampus and prefrontal cortex [46]. LDAM marker genes characterized in the context of Alzheimer’s disease [47], including *ACSL1*, *NAMPT*, *DPYD*, and *CD163*, are all significantly upregulated in PWH microglia across regions (Fig. S9). These findings reveal a dynamic shift in microglia from a homeostatic state to multifaceted reactive phenotypes characterized by strong immune responses and dysfunctional metabolic profiles reminiscent of those seen in neurodegenerative conditions, all under chronic HIV infection.

Finally, we performed multi-variate feature selection to select the most predictive genes that can best classify PWH and CTR microglia cells. We optimized a logistic regression classifier with *l*_1_ regularization using a 5-fold cross-validation on the INS microglia (see Methods for details). With in total 642 genes selected, the optimized model achieves 0.990 balanced accuracy on the test data. We further ranked the genes based on their feature importance (Fig. 5h with top 20 genes highlighted). Many of the top-ranked genes are associated with microglial activation signaling pathways and their regulations, underscoring their critical roles in PWH microglia and how they differ from CTR microglia [34, 48, 49, 50]. Additionally, we applied the model trained on INS data to classify the PWH microglia from CTR microglia in the PFC and VST data. The balanced accuracy is 0.935 for the PFC data and 0.873 for the VST data. This further illustrates the similarity of the transcriptomic differences between PWH and CTR microglia across regions and confirms the cross-region similarity observed at the univariate level (Fig. S6).

### 2.4 Astrocytic reactivity and metabolic dysfunction in HIV

Astrocytes are responsible for maintaining homeostasis in the brain, and while the exact effects of HIV on astrocytes are not known, there is literature that suggests astrocytes become reactive in the presence of HIV [51]. Our initial transcriptomic analysis across all astrocytes revealed significant changes in gene expression, (Fig. 3b and c), including downregulation of regulatory and reparative genes such as *DIPKC1* and *GINS3* [52, 53], alongside upregulation of markers associated with reactive astrocytes in other neurological diseases such as *SERPINA3*, *OSMR*, and *CHI3L1* [54, 55, 56] (Fig. 6). Over-representation analysis of top GO biological process pathways revealed several pathways related to cilium movement, signal transduction, and hypoxia associated pathways to be upregulated in HIV. Interestingly, cilia have been associated with neurotoxic and reactive astrocytes [57]. Moreover, many pathways related to translation and metabolism were downregulated in HIV. One of the main roles of astrocytes in the brain is to support glucose metabolism in the brain, and downregulation of these pathways in people with HIV suggests that there may be metabolic dysregulation occurring as a result of HIV infection [58] (Fig. 6b).

**Figure 6:**
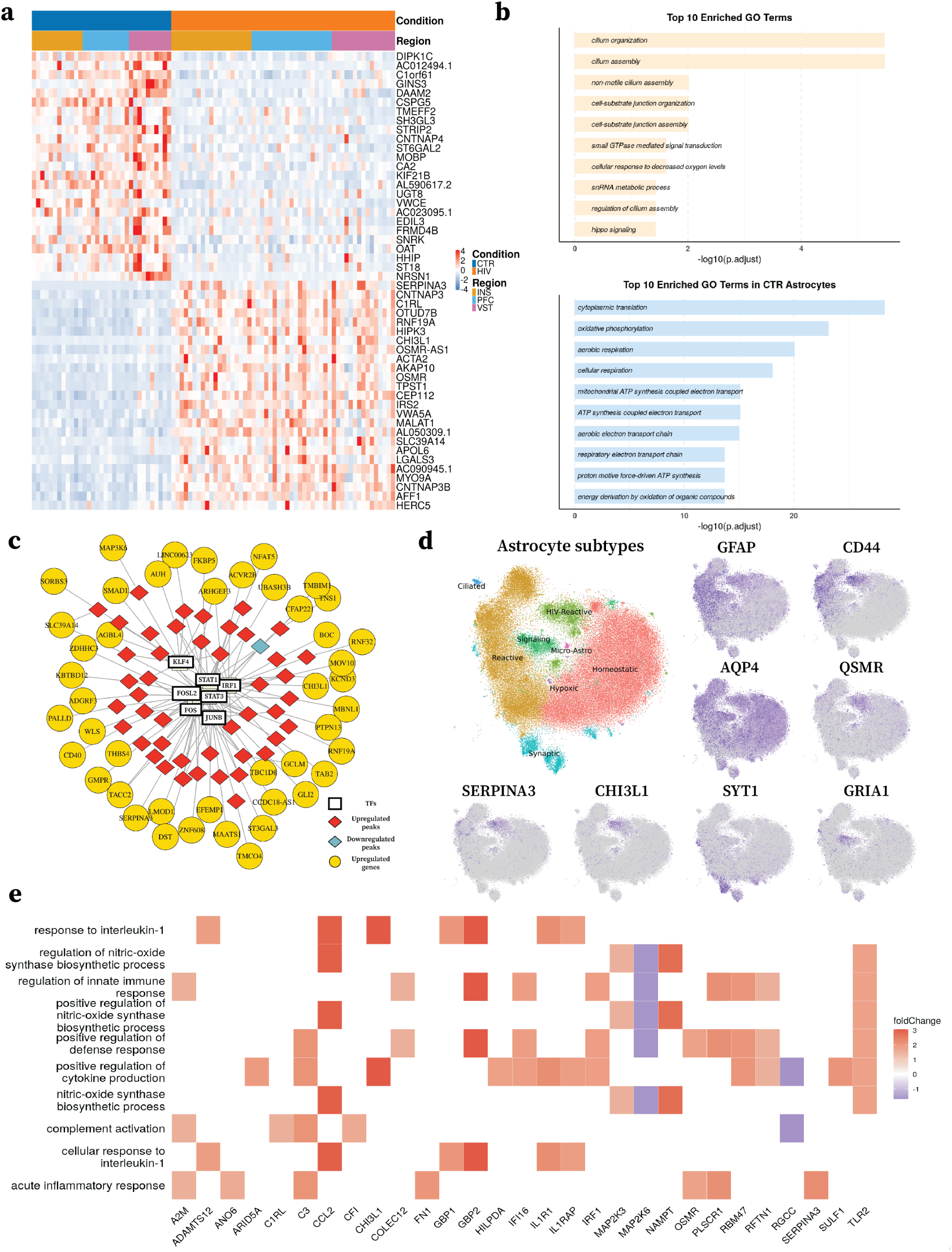
Translational and epigenetic changes on astrocytes of INS, PFC, VST due to HIV infection. **a**: Heatmap visualization of the top DEGs of the HIV main effect. Columns are different samples grouped by conditions and regions, and rows are the top DEGs with 25 each in the HIV-upregulated group and HIV-downregulated group. The expression levels inside the heatmap are z-scored for each gene. **b**: Top 10 enriched GO terms for the genes upregulated in PWH (top) and the genes downregulated in PWH (bottom) **c**: Gene regulatory network analysis conducted using MAGICAL for upregulated peaks and genes in PWH. A subset of transcription factors was chosen for visualization. The connections are visualized connecting transcription factors to peak and peaks to genes. **d**: t-SNE visualization of astrocyte subtypes and the expression levels of the corresponding marker genes. **e**: Heatmap of GO biological process based pathway analysis of markers of HIV-Reactive astrocytes colored based on log fold change of DEGs based on a wilcoxon test.

Motif analysis for each region further identified the enrichment of *JUN* -family transcription factors (TFs) in the HIV-upregulated peaks across brain regions, indicative of stress responses observed in other diseases, often accompanied by NF-*κ*B activation [59] (Fig. S10a). NF-*κ*B activation has been associated with pro-inflammatory responses to injury in astrocytes in other diseases [60]. Similarly, *FOS* -family TFs, known mediators of stress responses in the medial prefrontal cortex, were also enriched, suggesting a broader impact of HIV infection beyond specific brain regions [61]. We also detected the enrichment of *STAT3*, a key mediator of astrogliosis through the JAK/STAT pathway and is also related to the NF-*κ*B pathway [62]. Interestingly, we also see the enrichment of *EGR1* which has been shown to be upregulated due to HIV Tat1 in astrocytes [63], and *EGR4*, which has also been shown to be upregulated although not in astrocytes [64]. Collectively, these findings suggest that astrocytes in individuals with HIV adopt reactive phenotypes characterized by persistent inflammatory responses, even under antiretroviral therapy. Motif analysis of motifs enriched in the downregulated peaks for each region did not reveal any clear or consistent signatures across multiple brain regions (Fig. S10b).

Gene regulatory analysis of upregulated peaks and genes identified several networks involving reactive genes that were transcriptionally and epigenetically upregulated (Fig. 6c). Transcription factors such as *STAT3*, *STAT1*, and *FOS* exhibited increased regulatory activity, with associated upregulation of target genes. Notably, *SERPINA3* was linked to both *STAT3* and *FOS*, while *CHI3L1* was associated with *STAT1*, *IRF1*, and *FOSL2*. Additionally, *CD40*, a gene implicated in the production of inflammatory cytokines in astrocytes, was among the upregulated targets in these networks. This suggests a coordinated regulatory response driving astrocyte reactivity in the context of HIV infection and aligns with prior reports implicating *CD40* in inflammatory signaling in astrocytes [65].

Given the persistent reactive gene signature and the clear changes in metabolic pathways, we wanted to determine if these changes were due to specific subtypes of astrocytes or indicated broader astrocytic changes. To investigate astrocyte subtypes driving these changes, we analyzed 28,787 nuclei (7,910 CTR; 20,877 HIV) from the insula (INS), 20,627 nuclei (6,163 CTR; 14,464 HIV) from the prefrontal cortex (PFC), and 25,476 nuclei (7,398 CTR; 18,078 HIV) from the ventral striatum (VST). After excluding oligodendrocyte-contaminated clusters, astrocytes were significantly more abundant in HIV samples, particularly in the INS, as determined by a T-test(Fig. S11a), suggesting a potential expansion in astrocytes in PWH. Our analysis revealed several subpopulations, visualized using t-SNE (Fig. 6). Among these, we identified populations of *GFAP* +*CD44* + reactive astrocytes [66, 67] and a distinct *GFAP* +*CD44* +*OSMR*+*SERPINA3* + population of HIV-specific reactive astrocytes [55, 66, 67] (Fig. S11c). Additional subpopulations included *AQP4* +*GFAP* - homeostatic astrocytes [68], *SYT1* +*SNAP25* + synaptic astrocytes [69], and *GRIA1* + signaling astrocytes [70] (Fig. S11). Smaller and more patient-specific populations included *CFAP299* +*DTHD1* + ciliated astrocytes [71], *NR4A1-3* + hypoxic astrocytes [72], and *APBB1IP* + astrocytes expressing microglial genes [73] (Fig. S11b,c). These subtypes were defined based on gene expression within astrocyte-specific clusters, highlighting the functional diversity of astrocytes in HIV neuropathology.

We first examined the unique HIV-specific (*GFAP* +*CD44* +*OSMR*+*SERPINA3* +) reactive astrocyte subpopulation (HIV-Reactive) and found that the genes that mark that cell population, determined using a wilcoxon test, were involved in pathways related to defense response, interleukin signaling, complement activation, cytokine production, and nitric oxide synthase (Fig. 6e). Literature suggests that HIV activates the *C3* promoter through NF-*κ*B-mediated induction of *IL6*, contributing to HIV-mediated inflammation in patients with HAND [74]. Complement activation is also associated with synaptic loss and impaired neuronal function [74]. Although we did not observe increased expression of *IL1B* or *IL6*, HIV-Reactive displayed more abundant expression and elevated expression of *IL6R*, *IL1R*, and *C3*, suggesting that these phenomena may persist despite treatment with ART (Fig. S12). When examining ATAC-based gene activity in HIV-specific reactive astrocytes, we identified regional differences: in the INS and VST, pathways related to ECM reorganization and NF-*κ*B signaling were enriched. These results suggest that HIV-reactive astrocytes may contribute to the NF-*κ*B signature observed in RNA-based analyses (Fig. S12). This unique population of astrocytes enriched in HIV samples may contribute to the persistent pro-inflammatory signatures we see in astrocytes from people with HIV.

Changes in other astrocyte subtypes with abundant cell counts from both HIV and CTR samples were examined using a wilcoxon test to identify DEGs between PWH and CTR cells. Pathway analysis of DEGs upregulated in PWH showed signatures of response to virus and synaptic organization in several astrocyte subtypes Fig. S13. Pathway analysis of downregulated DEGs revealed consistent signatures of metabolic function or respiratory processes Fig. S13. This suggests that other astrocyte subpopulations are likely also being affected by HIV and may have altered function, and that there is a general trend of metabolic dysfunction of astrocytes in PWH.

Taken together, our analysis reveals clear and consistent reactive and pro-inflammatory changes in astrocytes and also suggests a decrease in metabolic function of astrocytes in the brains of people with HIV. These profound changes may be contributing to the prevalence of long-term consequences of HAND and other HIV-related disorders despite the use of ART therapy to manage HIV.

### 2.5 Brain Cell-Cell Communication and Interaction in HIV

Lastly, given the large amount of differential ligand/receptor signaling genes we identified in previous analyses, we performed cell-cell ligand-receptor analysis using NICHES [75], to examine the differential communication patterns across cell types between HIV and CTR (Fig. S14).

Between microglia and astrocytes, we observed strong interactions involving *SPP1* in INS, such as *SPP1* -*CD44* and *SPP1* -*ITGB1* (Fig. S14a). *SPP1* encodes a multifunctional phosphoprotein, and its expression level shows elevations under HIV infection and has been indicated to be associated with different neuro-degenerative diseases in microglia [76, 77, 78]. Its interaction with *CD44* and integrins such as *ITGB1* has been associated with a neuroinflammatory effect [79, 80]. These interactions appear to primarily occur in the INS based on our results. Another interesting interaction between microglia and astrocytes is *SYTL3* to *NRXN1*, which are both involved in synaptic function and may be contributing to potential synaptic alterations known to be associated with HIV, however, no direct link between these genes and HIV has been established [81, 82, 83]. These interactions suggest that microglia may be inducing reactivity in astrocytes in the brains of people with HIV and working together with astrocytes to alter synapses. These interactions appear to occur in both the INS and PFC and are notably absent from the VST, which has a different neuronal composition.

Astrocytes also appear to be interacting with microglia through complement related genes in primarily the INS. Astrocytes are known to use the complement system, particularly *C3*, to modulate astrocyte function and communicate with microglia and other cells [84]. We find several interactions between astrocytes and microglia related to *C3*, which has been linked to microglial activation in mice [85]. Moreover, we also see the *VCAN* -*TLR2* interaction, which may suggest microglial activation by astrocytes as *TLR2* activation in microglia has been associated with pro-inflammatory phenotypes and also is known to be upregulated in the presence of HIV [86, 34]. These interactions coupled with the microglial interactions to astrocytes suggest possible bidirectional communication between microglia and astrocytes where both cell types may be activating the other, but this would require subsequent biological experiments to confirm.

## 3 Discussion

Our understanding of the effects of HIV on the brain both regionally and at a cell type specific level is limited. The introduction of therapeutics has helped many PWH maintain suppression of viral replication and keep HIV encephalitis and severe forms of neurologic dysfunction at bay, but many PWH still face long term neurocognitive symptoms. In this study, we performed a comprehensive analysis of single nucleus multiomics on post-mortem brains from PWH decedents and healthy control decedents without HIV. Our analysis showed profound transcriptomic and epigenetic changes occur in association with HIV infection in multiple cell types and brain regions and helps highlight potential targets for therapeutics to mitigate some of the lasting effects of HIV on the brain.

PWH on ART often stop taking these medications towards the end of their lives, so it likely that some of the activation/reactive signatures that we see are a result of increased HIV replication close to death. If so, these findings nevertheless support that HIV is associated with detrimental effects and/or can recrudesce in the brain after years of therapy.

For the cell type annotations, we adopted a reference-based mapping strategy. INS annotations were derived primarily from human prefrontal cortex (PFC) references, while ventral striatum (VST) annotations relied on cross-species mapping using non-human primate atlases – a limitation stemming from the sparse availability of high-resolution, region-specific human brain cell atlases. Despite these limitations, conserved marker gene expression validates the robustness of our reference mapping approach for the known cell types.

There is ongoing discourse over the presence of a viral reservoir that persists during ART in the brain in microglia, though recent studies have lent support to this assertion that microglia are thought to be reservoirs of HIV in the brain because they are actively infected by HIV and are a population of long-lived cells [29]. Several studies have suggested that microglia may be integrating HIV into their genome, allowing for the possibility of latent HIV infection and causing a possible fundamental change in microglia function with HIV infection. We attempted to map HIV reads and found a limited number of cells that contained HIV DNA and transcripts. However, we lacked autologous HIV sequences for each patient, which have been shown to improve the accuracy of HIV detection [87]. Our limited mapping may also be due to the limitations of single nucleus studies, which are unable to identify cytoplasmic HIV RNA.

The molecular changes we documented, particularly in inflammatory, dysfunctional metabolic pathways, and glial cell function, provide a foundation for understanding HIV’s long-term effects on the brain. These findings point to specific cellular and molecular targets that may be relevant for addressing persistent HIV-associated neurological complications despite a history of ART treatment.

## 4 Acknowledgment

We thank the tissue donors and their families, Deborah Mash, Jennifer Chiarella, John Satterlee, and the SCORCH Consortium. This study is supported by National Institutes of Health (NIH) grants UM1DA051410 and U01DA053628. The authors gratefully acknowledge the support from the the National NeuroHIV Tissue Consortium (NNTC), financially supported by NIH funding through the NIMH and NINDS Institutes (grant nos. U24MH100930, U24MH100928, U24MH100929, U24MH100931, and U24MH100925). The contents are the responsibility of the authors and do not represent the official view of the NNTC or NIH.

## 5 Data and Code Availability

Data presented in this study were produced as part of the Single Cell Opioid Response in the Context of HIV consortium (SCORCH:RRID:SCR 022600). Publicly accessible data are available at NEMO Archive (RRID:SCR 002001) under identifier nemo: x. Access to all protected raw data associated with this study is managed by dbGaP and can be requested at x with the identifier x. Processed data are deposited under x. Codes for the data processing and analysis will be deposited to https://github.com/KlugerLab/Y-SCORCH_manuscript.

## 6 Methods

### 6.1 Study design

The primary goal of this study was to define cellular diversity and disease-associated transcriptomic changes and open chromatin profiles, as well as uncover gene regulatory interactions in the brains of patients with and without HIV. We performed 10x Genomics Multiome sequencing with both gene expression and assay for transposase-accessible chromatin with sequencing (ATAC-seq) from the same nuclei using postmortem human brain tissues from three brain regions: the prefrontal cortex (PFC), insular cortex (INS) and ventral striatum (VST). This study included 20 individuals with HIV and 13 sex- and age-matched healthy controls. Samples from HIV and control brains were paired into disease versus control groups based on their sexes and ages for batches of experiments. Investigators were not blinded to disease conditions during experiments and assessment of results.

Brain specimens of the prefrontal cortex, insular cortex, and ventral striatum from postmortem HIV patients and age- and sex-matched non-diseased control individuals were provided by brain tissue biorepositories at Manhattan Brain Bank, UCLA NeuroAIDS Bank, Texas NeuroAIDS Research Center, and Nova Southeastern University. Study participants were assigned into disease or control groups based on their HIV clinical diagnosis as described previously. The dissected brain tissues were fresh frozen and stored at −80 ° C until used for sequencing and validation assays.

### 6.2 Brain nuclei isolation

Nuclei were isolated from postmortem human brain tissues as previously described with modifications. 25-50 mg of fresh frozen brain tissue was added into 3 mL of ice-cold lysis buffer consisting of 250 mM sucrose, 25 mM KCl, 5mM MgCl2, 20mM Tris-HCl (pH 7.5), protease inhibitors, RNase inhibitor (80U/ml), 1mM DTT, and 0.1% (v/v) Igepal CA-630. DTT, RNase inhibitor, protease inhibitors, and Igepal CA-630were added immediately before use. The suspension was transferred to Dounce tissue grinder and homogenized with loose and tight pestles, 30 cycles each, with constant pressure and without introduction of air. The homogenate was strained through 40 um tube top cell strainer (Corning #352340) which was pre-wetted with 1ml wash buffer: (250 mM sucrose (Sigma #S0389), 25 mM KCl (Sigma #60142), 5mM MgCl2 (Sigma #M1028), 20mM Tris-HCl (pH 7.5) (AmericanBio #AB14043; Sigma #T2413), protease inhibitors w/o EDTA (Roche #11836170001), RNase inhibitor (80U/ml) (Roche #03335402001), 1mM DTT (Sigma #43186)). Additional 4 mL of wash buffer was added to wash the strainer. Final 10 mL of solution was mixed with 10 mL of 50% Optiprep (Axis-Shield# 1114542) solution (50% iodixanol (v/v), 250 mM sucrose (Sigma #S0389), 25 mM KCl (Sigma #60142), 5mM MgCl2 (Sigma #M1028), 20mM Tris-HCl (pH 7.5) (AmericanBio #AB14043; Sigma #T2413), protease inhibitors w/o EDTA (Roche #11836170001), RNase inhibitor (80U/ml) (Roche #03335402001), 1mM DTT (Sigma #43186)) by inverting the tube 10x and carefully pipetted into 2 centrifuge tubes (Corning #430791). The tubes were centrifuged at 1000 g, for 30 min at 4 C on centrifuge (Eppendorf #5804R) and rotor (Eppendorf #S-4-72). Upon end of centrifugation, the supernatant was carefully and completely removed and total of 5 mL of resuspension buffer (250 mM sucrose (Sigma #S0389), 25 mM KCl (Sigma #60142), 5mM MgCl2 (Sigma #M1028), 20mM Tris-HCl (pH 7.5) (AmericanBio #AB14043; Sigma #T2413), protease inhibitors w/o EDTA (Roche #11836170001), RNase inhibitor (80U/ml) (Roche #03335402001), 1mM DTT (Sigma #43186)) was added carefully on the pellets in tubes and centrifuged at 1000 g, for 10 min at 4 C on the same centrifuge and rotor. Supernatants were then carefully and completely removed, pellets were gently dissolved by adding 100 ul of resuspension buffer (see above) and pipetting 30x with 1ml pipette tip, pooled and filtered through 35 um tube top cell strainer (Corning #352340). Use the nuclei with 10x Genomics Chromium Next GEM Single Cell Multiome ATAC + Gene Expression as soon as possible with recovery target of 10,000 nuclei, using 5ul of final nuclei solution.

### 6.3 Single nucleus ATAC and RNA sequencing

The single nucleus ATAC and RNA sequencing libraries were prepared using the Chromium Next GEM Single Cell Multiome ATAC + Gene Expression chemistry, following the manufacturer’s instructions (10x Genomics). The generated single nucleus libraries were sequenced using Illumina NovaSeq6000 S4 sequencer at a sequencing depth of 300 million reads per sample, for both ATAC and gene expression libraries.

### 6.4 Data processing

We processed the raw RNA and ATAC fastq files using 10x Genomics cellranger-arc v2.0.2 with human reference (GRCh38) as the reference genome for alignment.

#### 6.4.1 RNA processing

For the RNA outputs from cellranger-arc, we establish an automated processing pipeline for each sample with Nextflow [7]:

##### Quality control

We first exclude X/Y chromosomes and mitochondrial genes to avoid nuclei clustering based on sex or nuclei quality following the protocol from [88]. Nuclei that express less than 500 genes or more than 7500 genes are filtered out. We use scDblFinder [89] to remove possible doublets with default parameters. Finally, we remove clusters from each sample that have high percentage of UMIs that map to mitochondrial genes (*>* 10%).

##### Cell annotation

For each sample, we map the processed RNA data to the reference data from multiple reference single-cell RNA-seq data [10, 11, 12]. We apply `scTransform` from Seurat v5 with default parameters to normalize each reference dataset and each sample from our data. We then apply PCA to each and use `FindTransferAnchors` and `TransferData` from Seurat to annotate cells from each sample of our data. Specifically, we annotate our INS and PFC samples using references from [10, 11]. We remove cells that have low prediction scores from both references (prediction.score *<* 0.5), or inconsistent coarse-grained cell-type annotations (Ext, Inh, and Non-neuronal cells) between the two sets. We then follow the annotation procedure from [13] to reconcile the two sets of annotations into a single set of cell annotations, where neuronal cells (Ext and Inh) are annotated based on the annotations from [10] and non-neuronal cells are annotated based on annotations from [11]. We then integrate all samples within each region using `IntegrateLayers` from Seurat with Harmony [90], then remove the clusters biased towards one sample.

For our VST samples, we used an additional reference [12]. We first integrate all samples from VST, then cluster data and examine marker genes of different neuron types (Ext, Inh, MSNs). We removed seven samples that are enriched with abundant Ext which might be introduced by sample dissection bias, and we further removed the remaining clusters that are enriched with Ext. We also remove cells that have low prediction scores from all three references (prediction.score *<* 0.5) and keep only cells that have consistent coarse-grained cell-type annotations (Inh, MSNs, and Non-neuronal cells) across references and manual annotations. We then annotate the Inh cells using annotations from [10] and the non-neuronal cells using annotations from [11]. For cells that are annotated as MSNs, we further define their annotations by mapping them to a refined MSN subset reference from [12].

After annotating cell types for each region separately, we further refined the non-neuronal cell annotations by integrating all non-neuronal cells together from all regions. We cluster cells in the integrated space and remove ambiguous clusters by manually examining marker gene expressions. We further label the macrophage clusters by examining marker genes COLEC12 and F13A1.

##### Cell annotation of astrocyte subtypes

Cell annotations of astrocytes subtypes was completed using canonical marker genes from literature and other astrocytes based studies. A cluster-based approach was used for annotation.

#### 6.4.2 ATAC processing

For the ATAC outputs from cellranger-arc, we processed the data using signac [9].

##### Quality control

For each sample in the ATAC modality, we retained only the nuclei that also passed the RNA QC steps and further filtered the nuclei by retaining nuclei that have (1) the total number of fragment counts for the nuclei within (1500, 100000), (2) nucleosome signal *<* 2, and (3) TSS enrichment score *>* 1.

##### Peak calling and gene activity

For the filtered data, we re-called peaks for each cell type in each region using MACS2 [91] as annotated by the RNA modality. The gene activity score is computed using signac by aggregating the counts within the gene body and 2000 bases upstream of each gene.

### 6.5 Differential analysis

#### Differential gene expression analysis

The differential gene expression analysis is performed using EdgeR [14] from Muscat [15]. Age, sex, race, and PMI are included as covariates in the design matrix in addition to the condition labels. We first performed the differential analysis for each cell type in each region separately. In addition, we performed differential analysis across all regions to examine the average effect of HIV. Specifically, we built the contrast (INS.HIV-INS.CTR)/3+(PFC.HIV-PFC.CTR)/3+(VST.HIV-VST.CTR)/3 in EdgeR. P-values are adjusted for multiple testing across genes and cell types [15]. We first filter the DEGs based on the percentage of expressions: we compute the percentage of cells that express each gene in each sample and only retain genes that have an average percentage of expression *>* 0.05 across the HIV samples or across the CTR samples. We then filter the DEGs based on their log fold change and adjust p values: abs(logFC) *>* 0.25 and adjust p values *<* 0.05.

#### Differential ATAC analysis

Similar to differential gene expression analysis, we performed differential peak analysis and differential gene activity analysis using EdgeR, with age, sex, race, and PMI included as covariates in the design matrix. The differential peaks and active genes are filtered with the same cutoffs as in the filtering steps of the differentially expressed genes.

#### Differential Expression Analysis of Astrocyte Subtypes

Differential expression analysis of astrocyte subtypes was conducted using a wilcoxon rank sum test using FindMarkers due to sample variability and more limited cell counts.

### 6.6 Regional & cross-region pathway analysis

For the identified DEGs, we performed Gene Ontology (GO) over-representation analysis using clusterprofiler [92]. We focus on biological process (BP) as the main ontology and report p values adjusted by the Benjamini-Hochberg (BH) method. For the regional DE analysis, we filtered the pathways for each cell type in each region by retaining the top five pathways with the smallest p values and having at least three genes enriched in the pathway. For the cross-region analysis, we filtered the pathways by the adjusted p values *<* 0.05.

### 6.7 Gene correlation analysis

To identify correlated gene modules in HIV microglia, we first select DE genes using EdgeR [14, 15]. DE genes were filtered based on the following criteria: logFC *>* 1, adjusted p values *<* 0.01, logCPM *>* 2. Next, we refined the cell population by removing clusters dominated by a single sample. Cells were clustered using the FindClusters function in Seurat with resolution = 1.5. Clusters where a single sample contributed more than 50% of the cells were excluded, and the remaining cells were retained for correlation analysis.

For the selected genes and filtered cells, we computed gene-gene pairwise correlations using the CS-CORE R package [41]. Significantly correlated gene pairs (BH-adjusted p values *<* 0.05) were preserved for downstream analysis. Gene modules in HIV microglia were identified using hierarchical clustering with average linkage, followed by careful examination of the dendrogram to define modules containing at least 40 genes [93]. For each identified module, correlations were calculated in CTR microglia to assess whether the module structure was preserved.

### 6.8 Multi-variate feature selection

We first log-normalized the processed counts of RNA microglia, then randomly split the HIV/CTR samples into a training set and a test set. We built a logistic regression classifier with *l*_1_ regularization to classify the INS HIV microglia from INS CTR microglia using their genes as features. Within the training set, we optimize the model using a 5-fold cross-validation scheme. Specifically, we first split the training HIV/CTR samples into 5 groups where each group has randomly assigned HIV and CTR samples. We then evaluate a grid of the regularization parameter *λ* from [1*e^−^*^3^, 10]. For each *λ* in the grid, we train the corresponding model on 4 groups of the HIV/CTR samples and validate the balanced classification accuracy on the remaining group (validation set). We select the *λ* that gives the largest accuracy on the validation set and re-train the model using all 5 groups within the training set, and test its performance on the left-out test set. Lastly, We tested the model performance trained on INS microglia on the PFC data and VST data.

### 6.9 Motif analysis

We use JASPAR2020 [94] as the motif database and FindMotifs from signac to identify motifs that are over-represented in the differential peaks for each cell type in each region. We filter the motifs by adjusted p-values *<* 0.05 and compute motif activities using ChromVAR [95]. To visualize the motif activities, we computed the average motif activities grouped by condition in each region and combined the results across regions for each cell type.

### 6.10 Gene regulatory network analysis

We used the Bayesian framework MAGICAL [96] to infer the gene regulatory network in astrocytes. Genes upregulated in HIV (adjusted p values *<* 0.05 and abs(logFC) *≥* 0.5) are selected as candidate genes, and peaks more accessible in HIV with the same thresholds are selected as candidate peaks. Genes were annotated using the GRCh38 reference genome, and transcription factor binding motifs were identified using the JASPAR2020 [94]. The analysis considered potential regulatory linkages between peaks and genes within 500 kb of each other. The regulatory network was visualized using Cytoscape [97] and igraph [98], focusing on selected genes of interest.

### 6.11 Cell communication analysis

We ran NICHES [75] for each sample to analyze the cell-cell communication patterns between cell types. We only include samples that have more than 500 cells as input to NICHES, then we concatenate the output CellToCell ligand-receptor matrix from NICHES across samples. For each cell type pair (e.g., Astro-Micro), we performed differential ligand-receptor analysis between HIV and CTR using FindMarkers from Seurat. We filtered the interactions by adjusted p-values *<* 0.01 and reported the top 20 interactions ranked by log fold-change for each condition.

### 6.12 Viral detection

#### HIV Transcripts

A collection of HIV-1 clade B transcripts was collected from the HIV Sequence Database [99]. BLAST was then used to align these sequences with the human genome GRCh38, and any transcript that produced an alignment was removed from the collection. HXB2 was then added to this list to complete the list of HIV transcripts utilized for all further HIV detection.

#### RNA Detection

We built a pipeline to detect viral RNA based on the work published by Wei et al. [100] using STAR [101]. In addition to their methods, we also removed all reads containing erroneous polyG sequences of 15 or longer. We then collapsed identical reads that may be due to PCR amplification and counted the number of reads per cell whose barcode matched or was within a Hamming distance of the 10X Multiome whitelist. A threshold was used such that any cell barcode with *>* 2 HIV-aligning reads was considered to contain HIV RNA.

#### DNA Detection

To identify viral DNA reads, we built a pipeline based on the work published by Wei et al. [100] using bowtie2 [102]. In addition to their methods, we also removed all reads containing erroneous polyN sequences of 15 or longer. We then collapsed identical paired reads that may be due to PCR amplification and counted the number of reads per cell whose barcode matched or was within amming distance of the 10X Multiome whitelist. We also removed all paired reads that aligned with the human genome GRCh38 using bowtie2 with the same settings as those used to align the reads with our collection of viral transcripts. A threshold was used such that any cell barcode with *>* 2 HIV alignment reads was considered to contain HIV DNA.

## S1 Supplementary Figures

**Figure S1:**
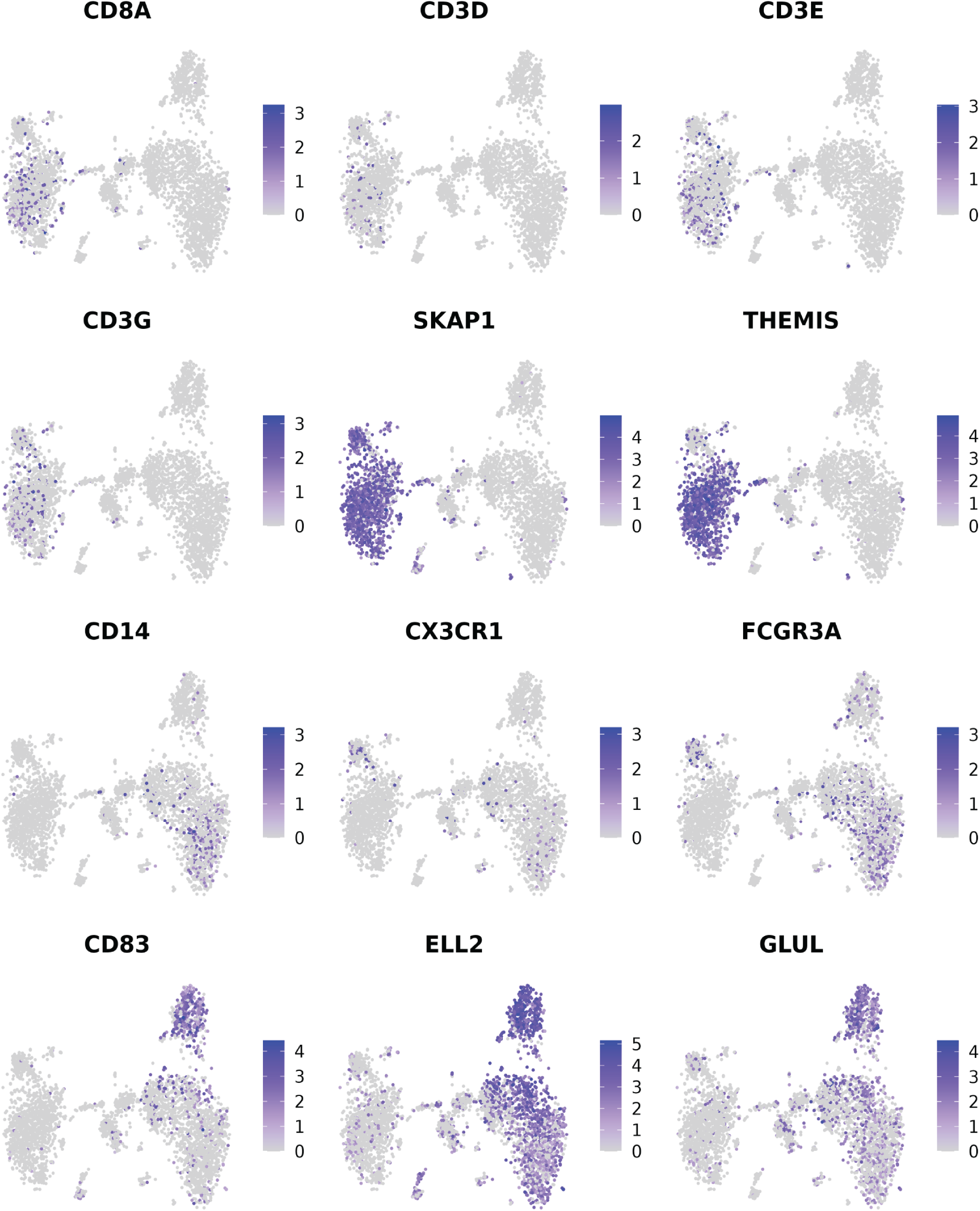
t-SNE visualization of immune marker genes of the identified immune population.

**Figure S2:**
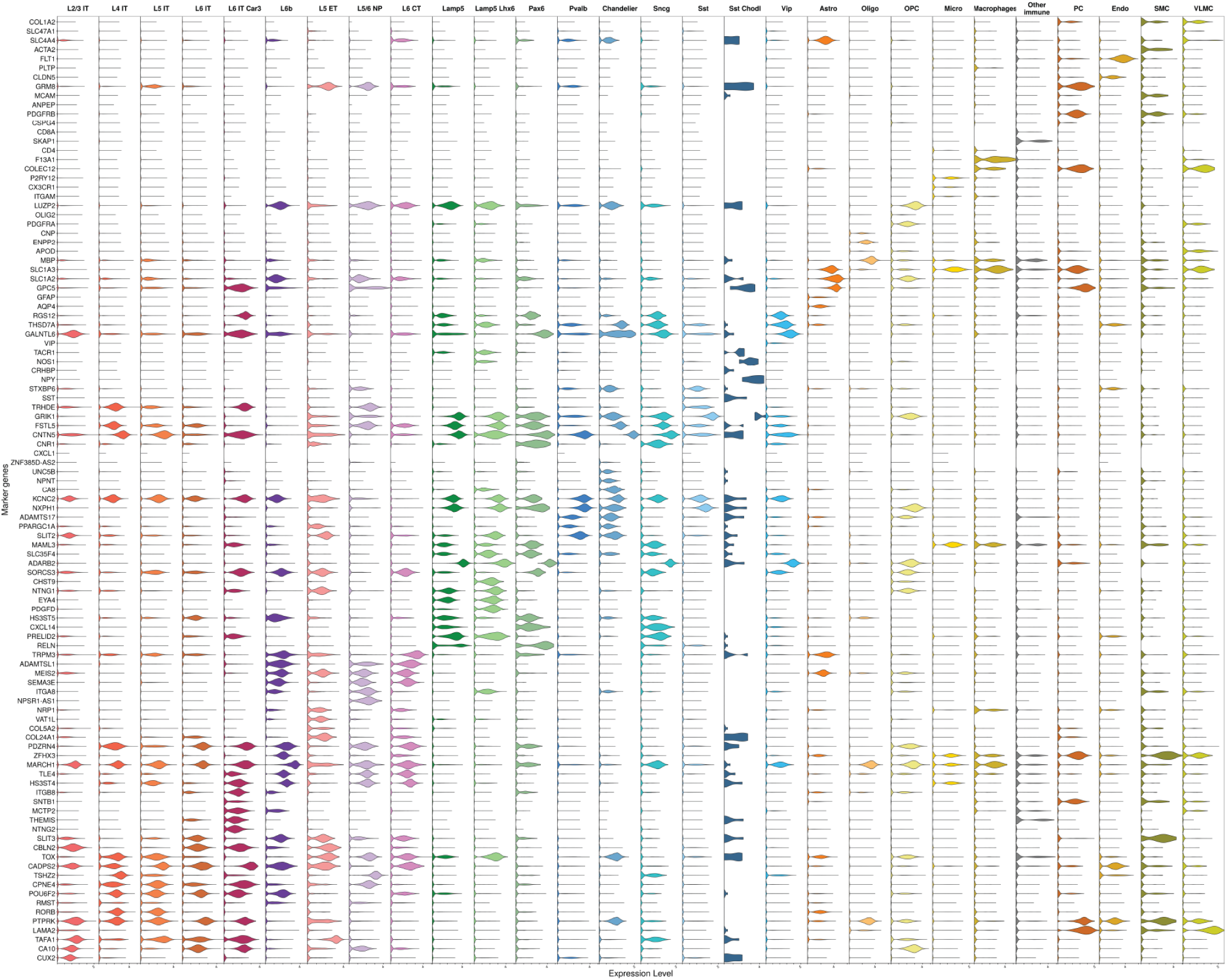
Visualization of the expressions levels of marker gene (y-axis) across all cell types (x-axis) in PFC.

**Figure S3:**
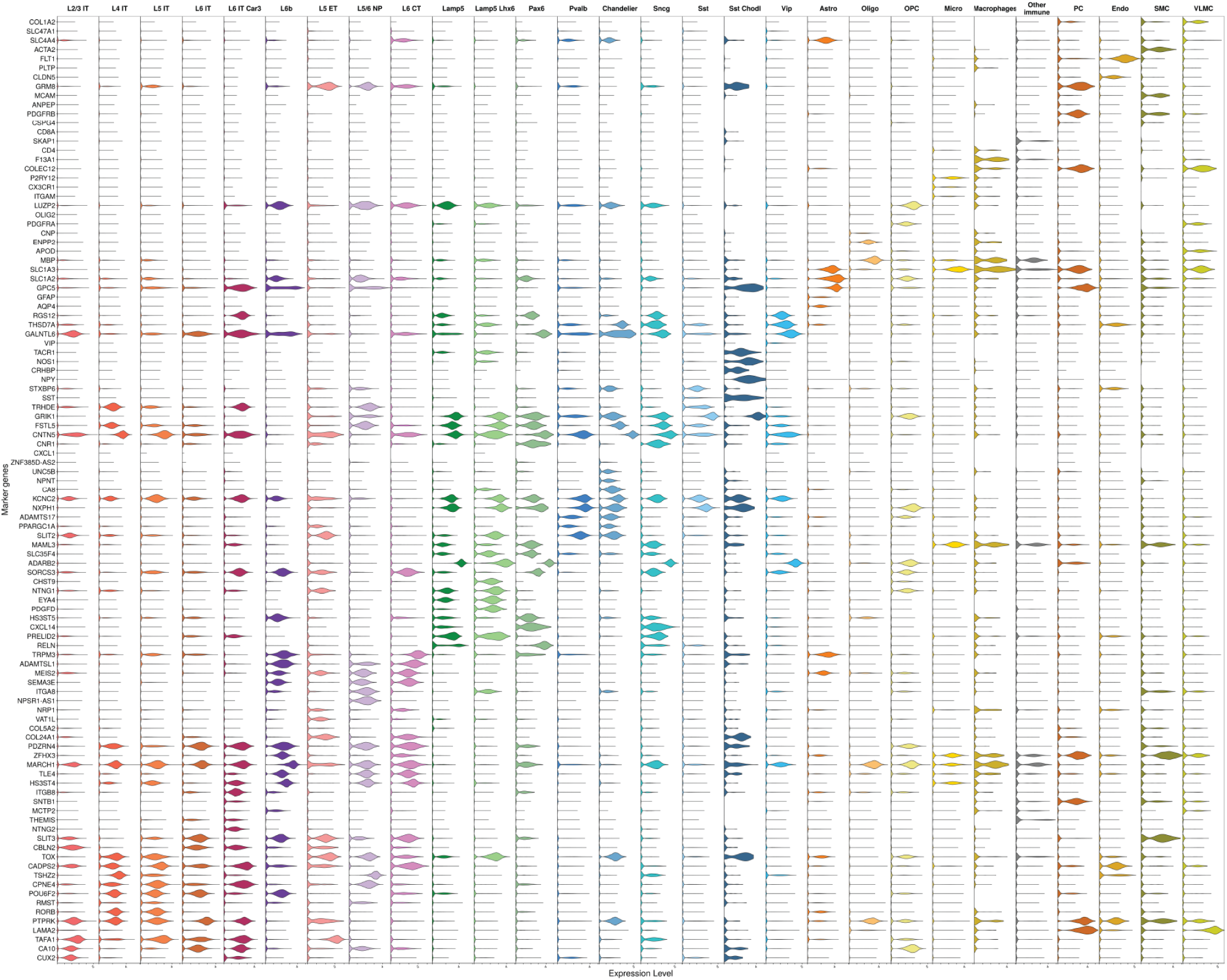
Visualization of the expressions levels of marker gene (y-axis) across all cell types (x-axis) in INS.

**Figure S4:**
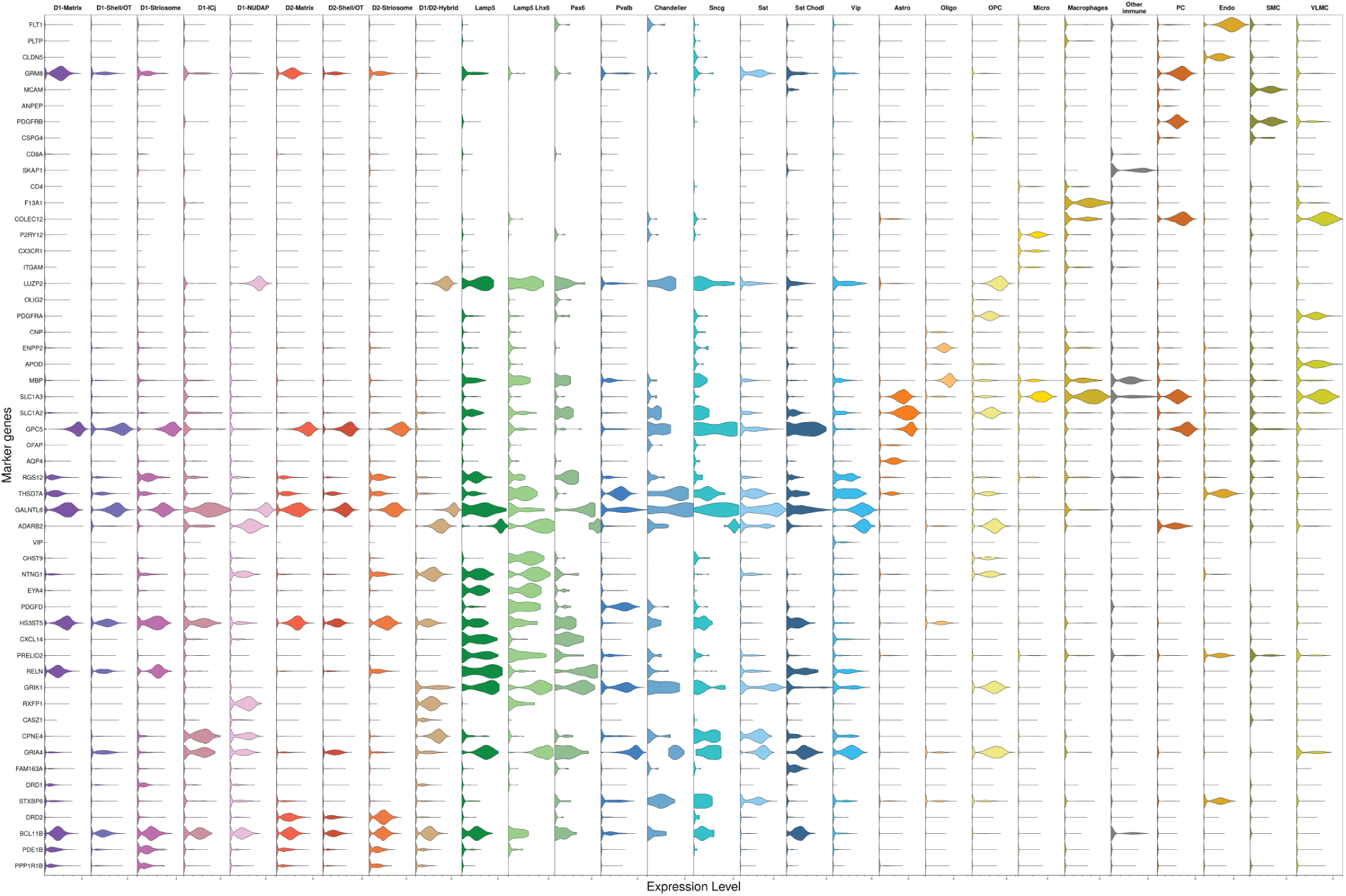
Visualization of the expressions levels of marker gene (y-axis) across all cell types (x-axis) in VST.

**Figure S5:**
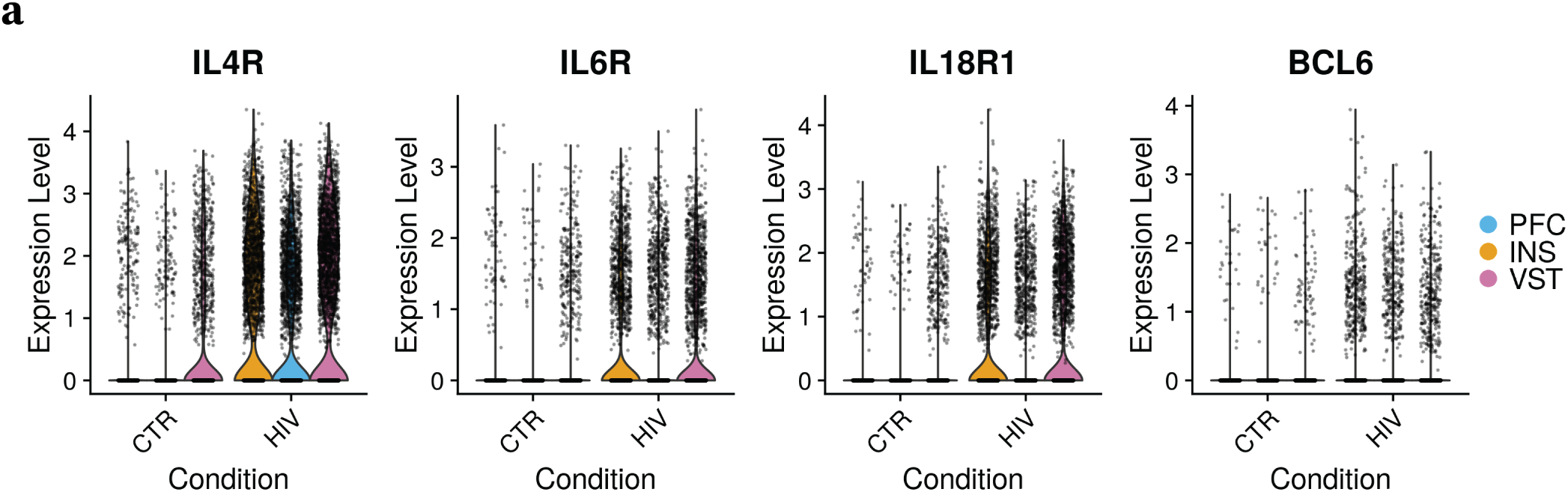
Comparison of expression levels of several cytokine receptor genes between HIV and CTR in endothelial cells across regions. The violin plots in each condition is split by regions (INS, PFC, VST, from left to right).

**Figure S6:**
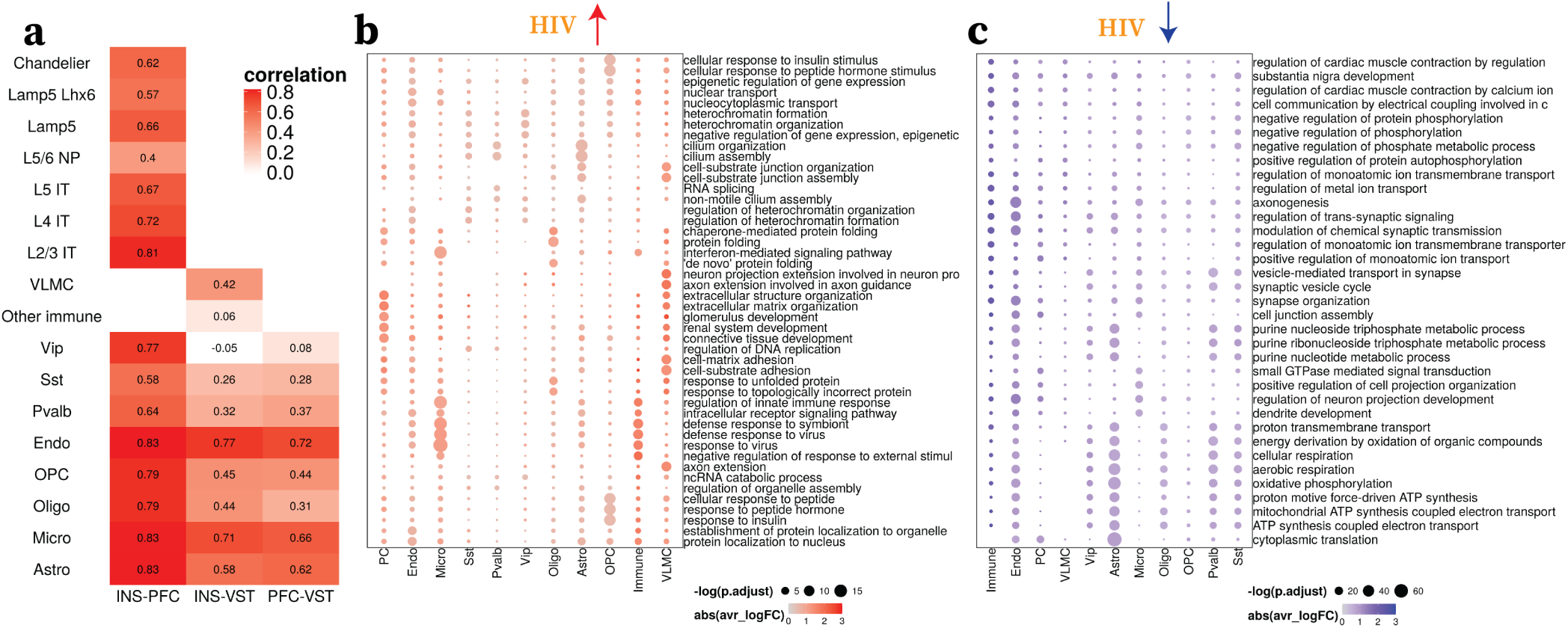
Cross-region comparison of differential signals. **a**: Heatmap visualization of Spearman correlation coefficient of logFC across DEGs between each pair of regions (columns) for each cell type (rows). **b**: Top pathways (rows) upregulated in HIV for the average HIV effect across all regions (columns). **c**: Top pathways (rows) downregulated in HIV for the average HIV effect across all regions (columns).

**Figure S7:**
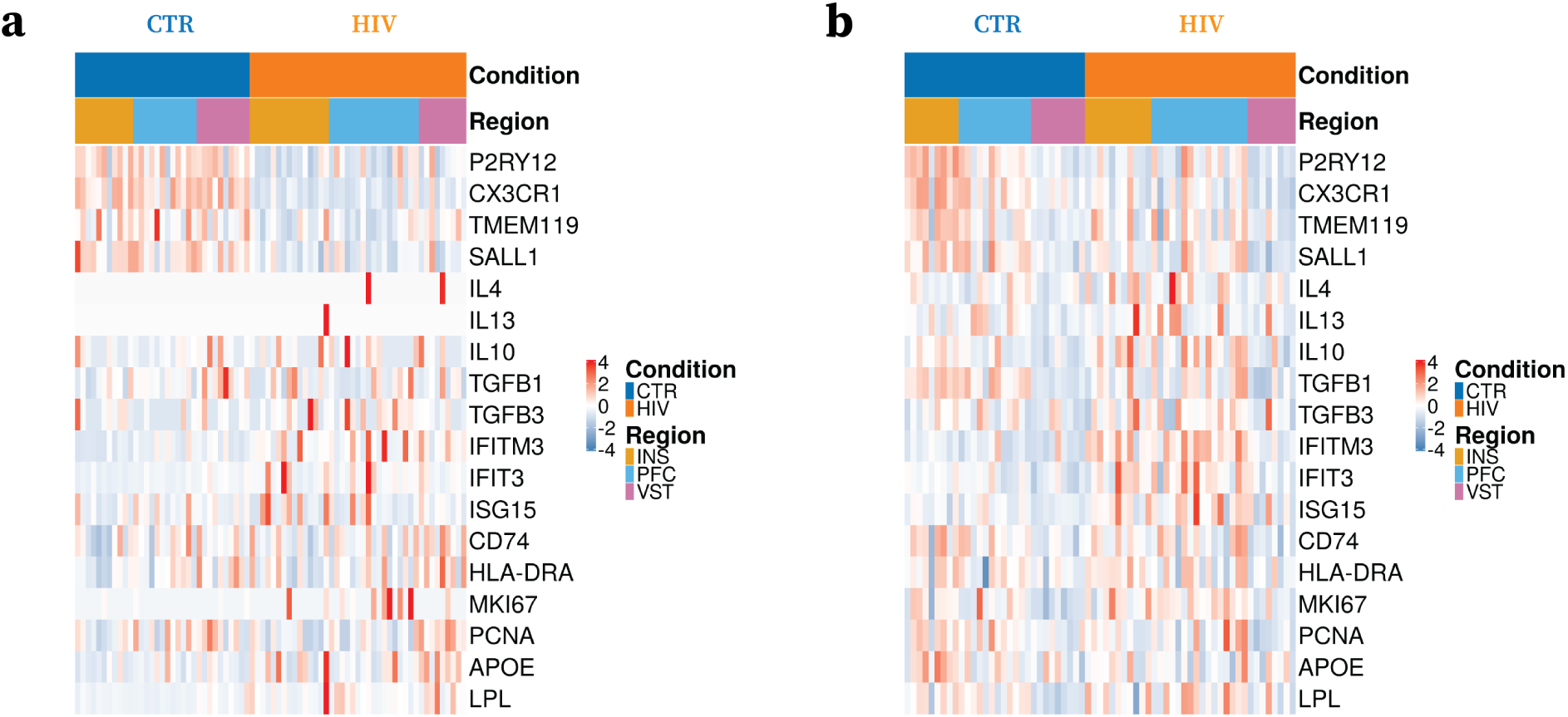
Heatmap visualization of different microglia state markers across samples. **a**: Comparison of gene expression levels **b**: Comparison of gene activities. Columns are samples grouped by conditions and regions, and rows are marker genes. Homeostatic markers: P2RY12, CX3CR1, TMEM119, SALL1; Anti-inflammatory markers: IL4, IL13, IL10, TGFB1, TGFB3; Interferon-responsive markers: IFITM3, IFIT3, ISG15; Antigen presentation markers: CD74, HLA-DRA; Proliferative markers: MKI67, PCNA; Disease–associated markers: APOE, LPL.

**Figure S8:**
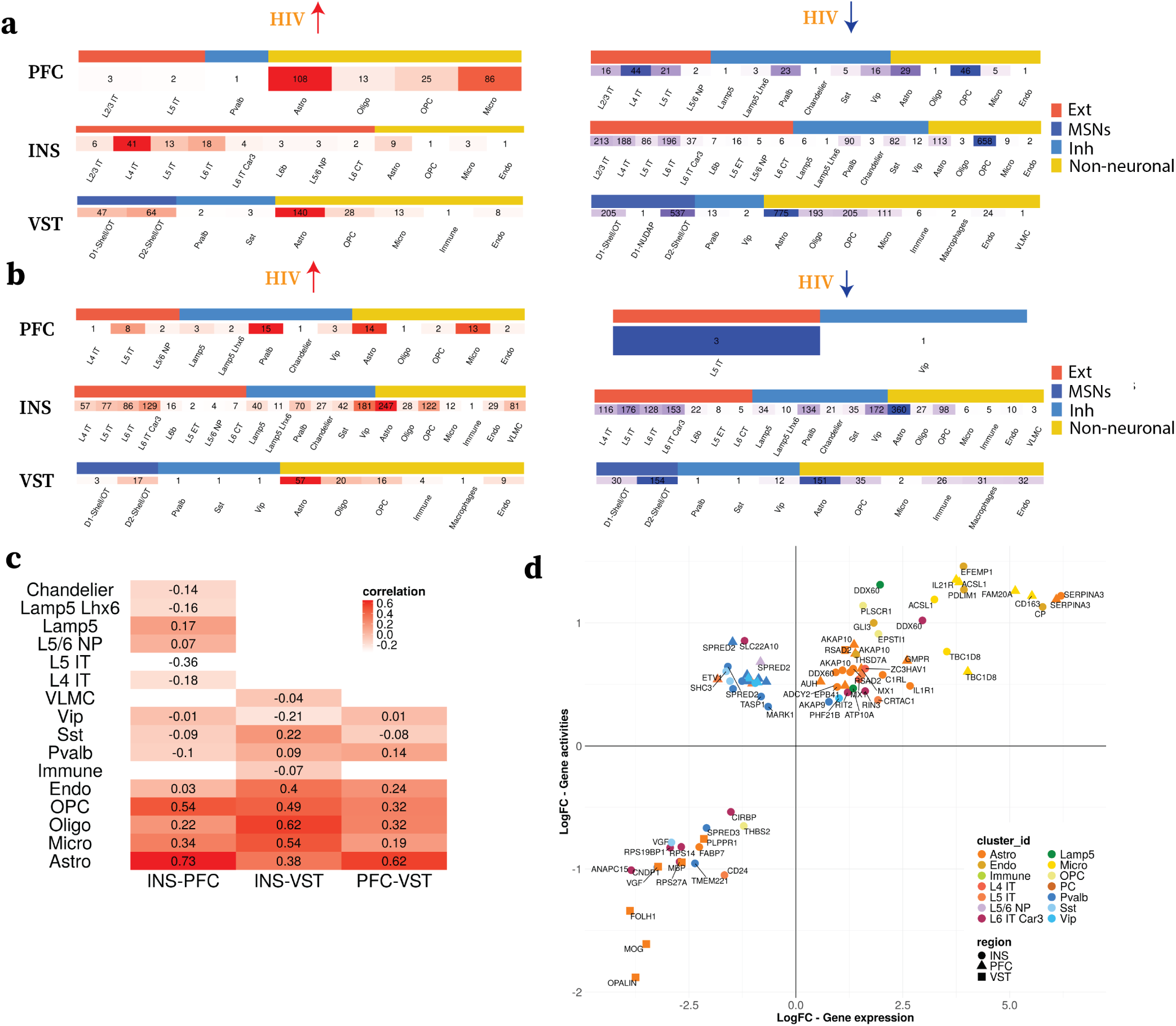
Epigenetic differential analysis reveals strong HIV effect. **a**: Total number of differential peaks for each cell type that are upregulated in HIV (left) and downregulated in HIV (right). The heatmap is splited by regions and colored by coarse-grained cell types (Ext, MSNs, Inh, Glial). **b**: Total number of differential peaks for each cell type that are upregulated in HIV (left) and downregulated in HIV (right). The heatmap is splited by regions and colored by coarse-grained cell types (Ext, MSNs, Inh, Glial). **c**: Heatmap visualization of Pearson correlation coefficient of logFC across differential gene activities between each pair of regions (columns) for each cell type (rows). **d**: Comparison of the effect size (logFC) between differential gene expressions (x-axis) and differential gene activities (y-axis) for top genes. Genes are colored by cell types. Regions are indicated by shapes.

**Figure S9:**
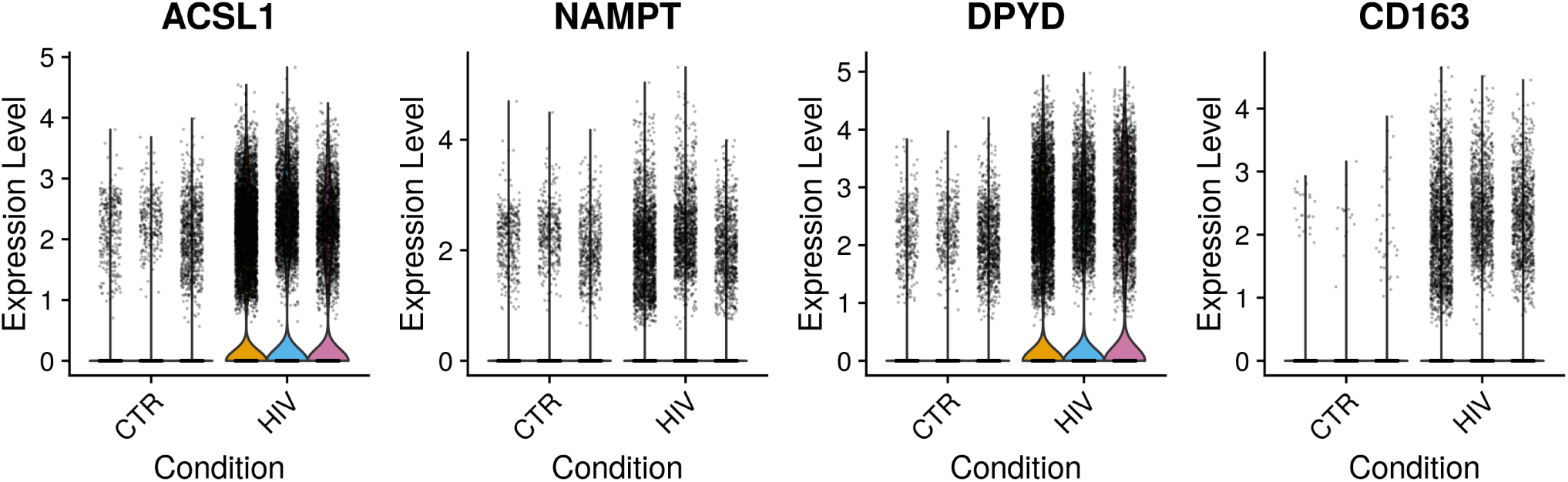
Comparison of expression levels of markers genes associated with Lipid droplet accumulating microglia (LDAM). The violin plots in each condition is split by regions (INS, PFC, VST, from left to right).

**Figure S10:**
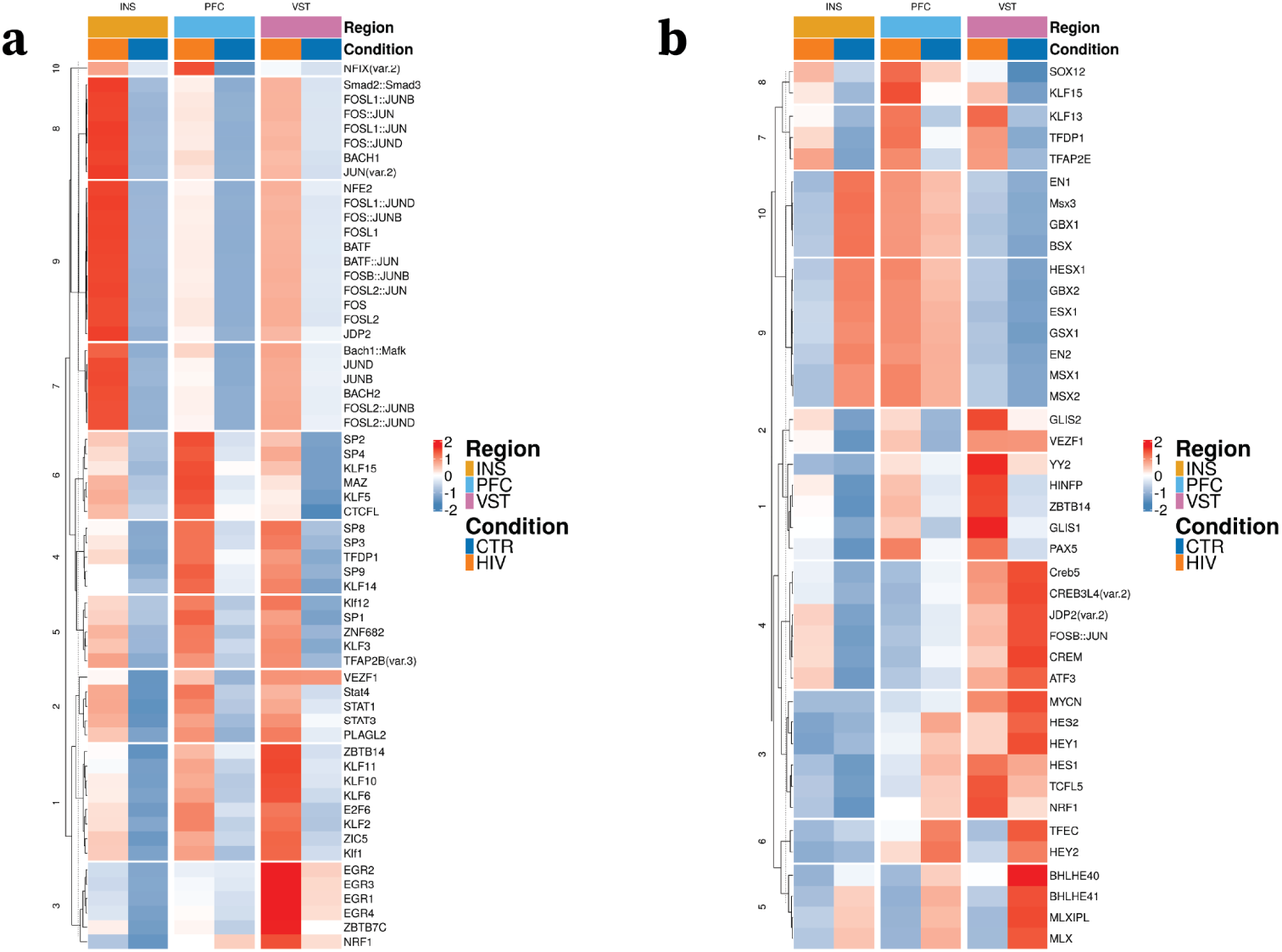
**a** Heatmap visualization of the activities of motifs enriched in HIV upregulated genes from astrocytes. The average motif activities by region and conditions are shown for each motif after z-scoring. Motifs are clustered by k-means clustering.**b** Heatmap visualization of the activities of motifs enriched in HIV downregulated genes from astrocytes. The average motif activities by region and conditions are shown for each motif after z-scoring. Motifs are clustered by k-means clustering.

**Figure S11:**
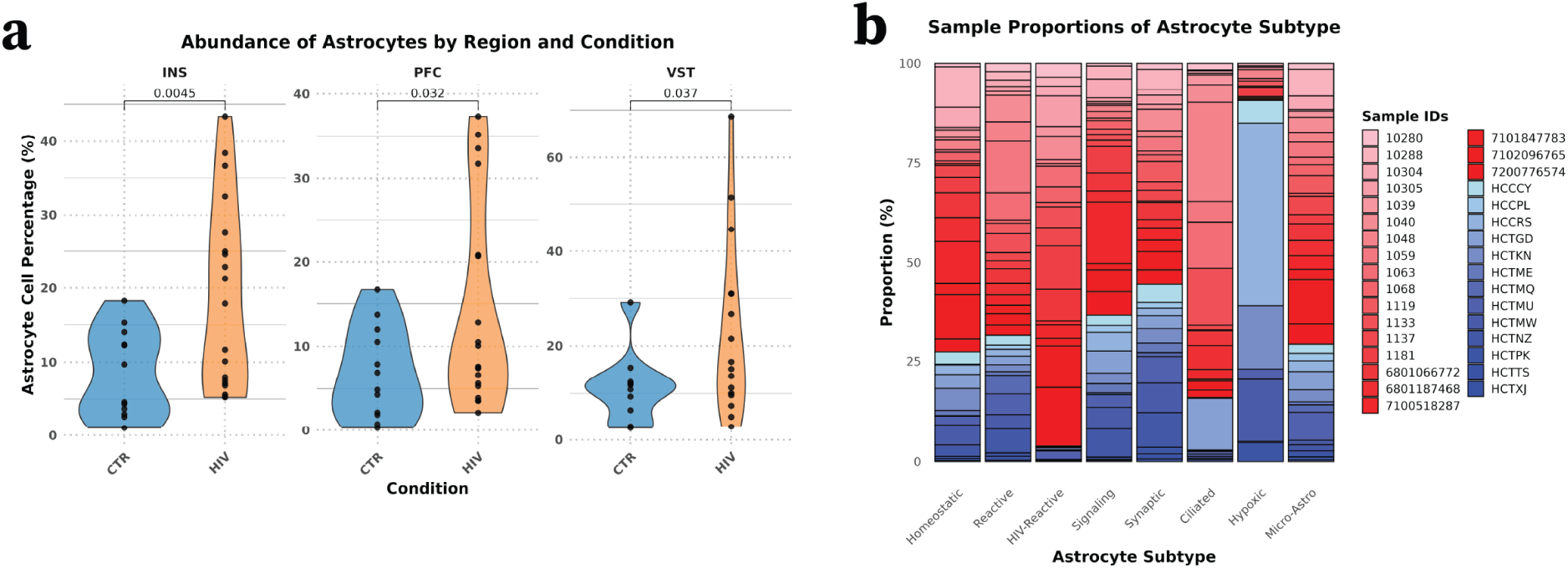
Astrocyte subtype supplemental figure. **a**: Violin plot of abundances of astrocytes from each sample for the INS, PFC, and VST. **b** Bar plot of sample proportions for astrocyte subtypes.

**Figure S12:**
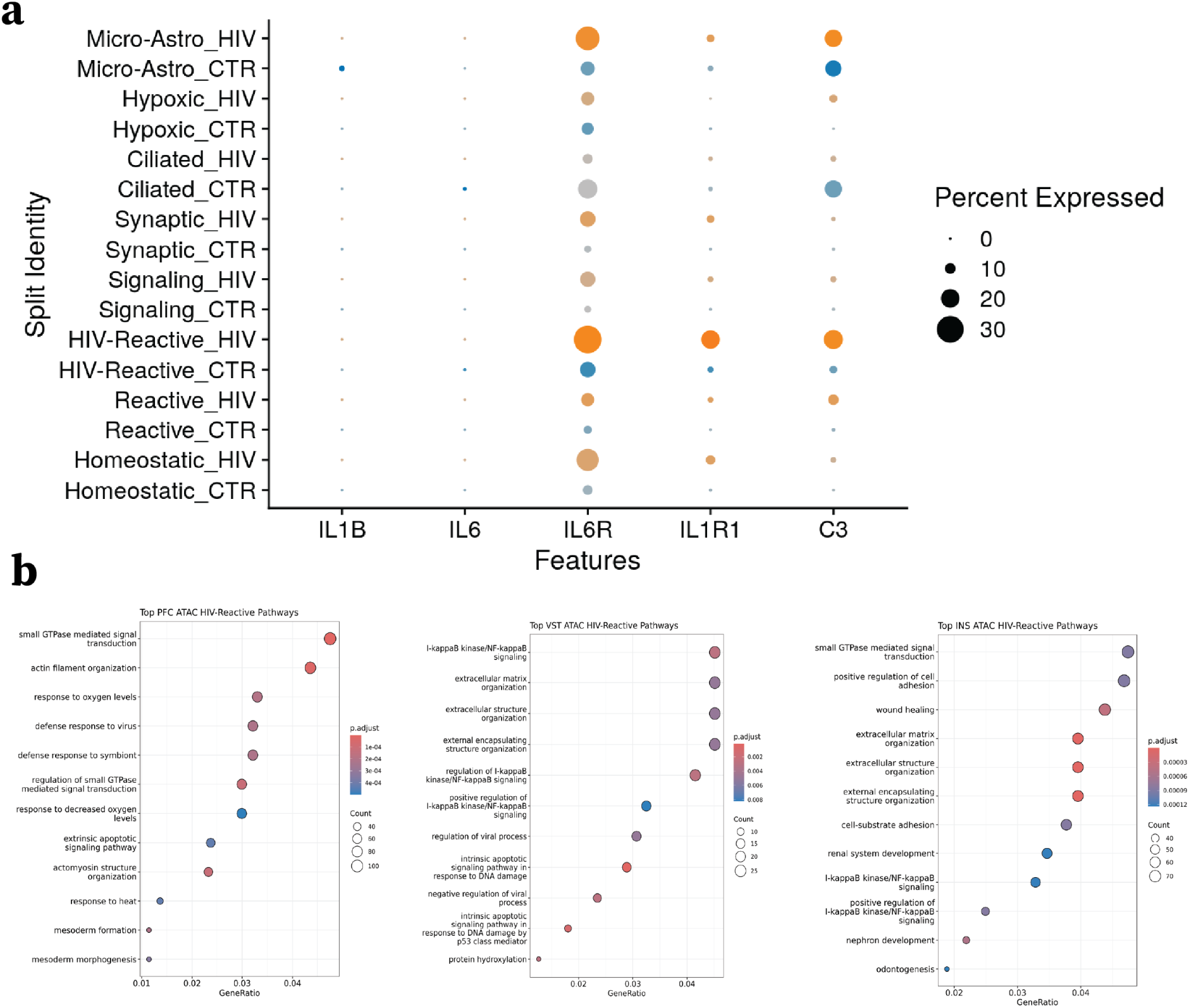
HIV Reactive ATAC pathway supplemental figure. **a**: Dotplot visualization of interleukin and complement genes expression comparing HIV vs CTR samples. **b** Pathway analysis of genes with top gene activity from snATAC data from the HIV-Reactive cluster of cells for the INS, PFC, and VST.

**Figure S13:**
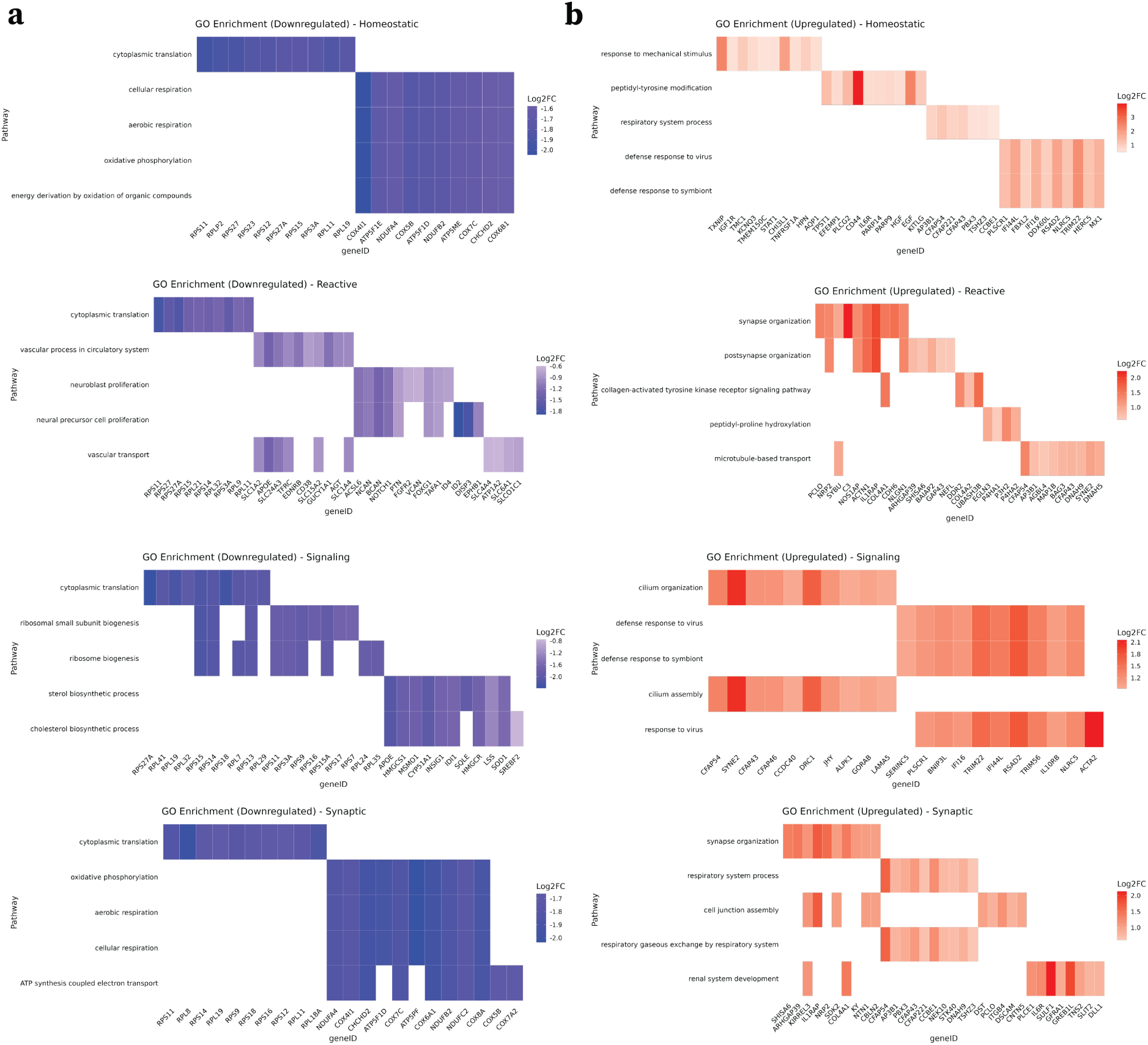
Pathway analysis for other astrocyte populations. **a**: Heatmap visualization of top 5 pathways colored by logFC of 10 genes present in those pathways from downregulated DEGs of homeostatic, reactive, signaling, and synaptic astrocytes. **b**: Heatmap visualization of top 5 pathways colored by logFC of 10 genes present in those pathways from upregulated DEGs of homeostatic, reactive, signaling, and synaptic astrocytes.

**Figure S14:**
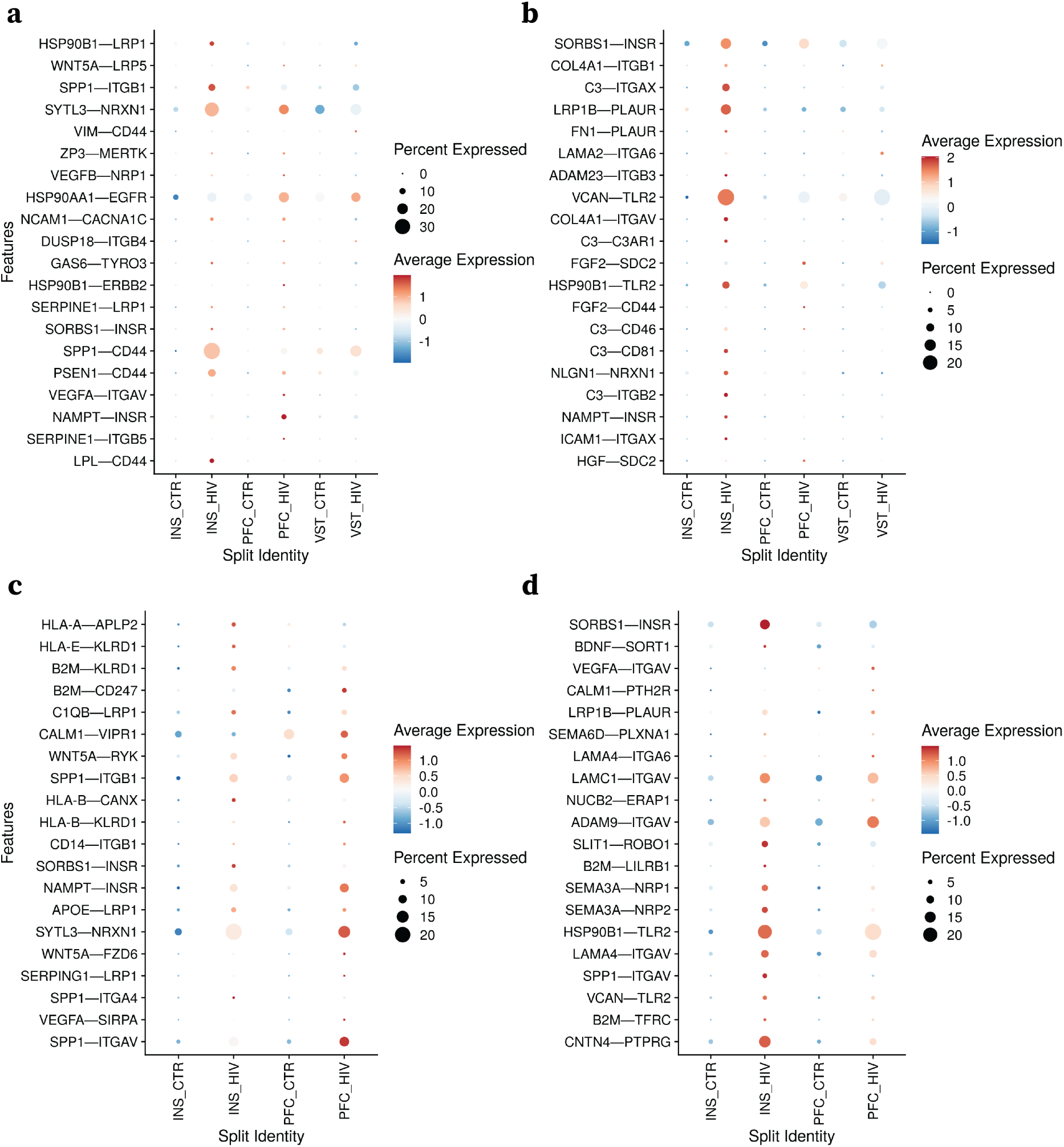
Dotplot visualization of the ligand-receptor interaction activities between cell types: **(a)** microglia - astrocytes, **(b)** astrocytes - microglia, **(c)**: microglia - L4 IT excitatory neurons, **(d)**: L4 IT excitatory neurons - microglia. For each panel, rows (y-axis) are the top 20 upregulated interactions in HIV, and the interactions are grouped by regions and conditions on the columns (x-axis).

